# Nanoscale curvature promotes high yield spontaneous formation of cell-mimetic giant vesicles

**DOI:** 10.1101/2020.07.29.227686

**Authors:** Joseph Pazzi, Anand Bala Subramaniam

## Abstract

To date, surface-assisted assembly of cell-like giant vesicles use planar surfaces and require the application of electric fields or dissolved molecules to obtain adequate yields. Here, we present the use of nanoscale surface curvature and hydrophilic surface chemistry to promote the high yield assembly of GUVs. We show that assembly on surfaces composed of entangled hydrophilic nanocellulose fibers results in an unprecedented 100,000-fold reduction in costs while increasing yields compared to extant techniques. Quantitative measurements of yields provide mechanistic insight on the effect of nanoscale curvature and the effect of surface chemistry. We present a thermodynamic ‘budding and merging’, BNM, model that unifies observations of assembly. The BNM model considers the change in free energy by balancing elastic, adhesion, and membrane edge energies in the formation of surface-attached spherical buds. Due to curvature and the hydrophilicity of cellulose, energetically unfavorable formation of buds on planar and spherical surfaces becomes favorable (spontaneous) on surfaces composed of cylindrical cellulose nanofibers.

**TOC Graphic:** 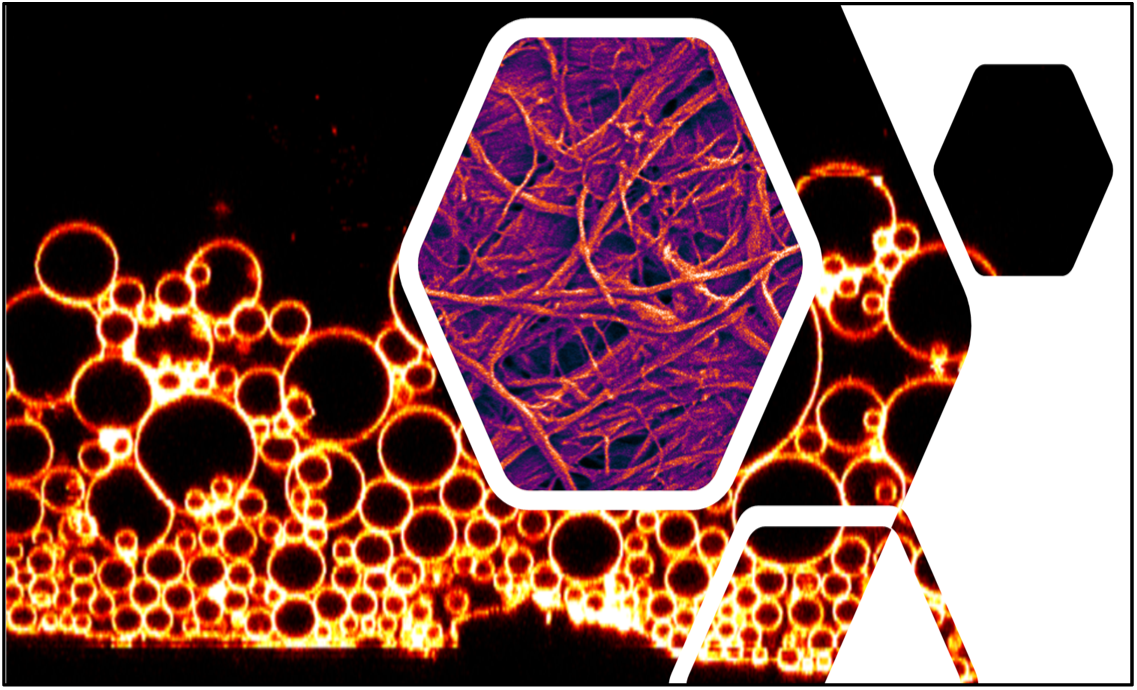

## Introduction

Giant unilamellar vesicles or GUVs are single-walled closed phospholipid bilayer membranes with diameters larger than 1 micrometer^1,2^. GUVs resemble minimal biological cells^1,2^. This resemblance has made GUVs instrumental in advancing understanding of, among others^3–7^, membrane organization^8–10^, cytoskeletal mechanics^11^, reproduction^12^, division^13–16^, transport^17,18^, and electrical signaling^19^. Furthermore, innovations in bottom-up synthetic biology^20–22^, engineering of artificial tissues^23,24^, and delivery of drugs^25,26^ are opening up new avenues for the use of GUVs in biomimetic applications.

Surface-assisted assembly, a growing collection^27–34^ of techniques in which solvent-free lipid films are templated onto surfaces and then hydrated in aqueous solutions, is a promising route to obtain giant vesicles. In the earliest discovered technique^27,28^, contemporarily known as ‘gentle hydration’^1,2^, dry lipid films are templated onto impermeable glass and Teflon surfaces and then hydrated in quiescent solutions. Electroformation modifies the procedure by applying an electric field normal to lipid films templated on conductive surfaces^29,30^. Other recent modifications include gel-assisted hydration, where lipid films are templated onto soluble hydrogels supported on glass surfaces^31,32,35^ and PAPYRUS, Paper-Abetted amPhiphile hYdRation in aqUeous Solutions, where lipid films are templated onto insoluble fibers of cellulose filter paper^33^ or fabrics composed of natural, semi-synthetic, and synthetic fibers^34^.

Herein we present the use of nanoscale surface curvature of hydrophilic cellulose nanofibers to assemble GUVs. Use of surfaces with nanoscale curvature results in significant procedural simplifications and a 100,000× fold reduction in the costs of assembling GUVs. We perform systematic experiments and show that lab-made nanocellulose paper and commercial tracing paper, both composed of entangled hydrophilic nanoscale cellulose fibers, significantly increases the yield of GUVs compared to growth on planar surfaces, including when compared to surfaces which are permeable to water and when compared to the widely-used electroformation technique. Further, we find that when the surfaces are rendered hydrophobic, the yield of vesicles is significantly reduced on both planar surfaces and on surfaces composed of nanoscale cylindrical fibers. To the best of our knowledge, this paper presents the first quantitative comparison of the yields of GUVs obtained from multiple surface-assisted assembly techniques.

To explain our results, we develop a thermodynamic <u>B</u>udding a<u>N</u>d <u>M</u>erging (BNM) model for the surface-assisted assembly of GUVs. The BNM model considers the free energy change of a membrane on a surface transitioning to a surface attached spherical bud. A balance between elastic, adhesion, and membrane edge energies shows that the transition requires the input of energy (endergonic) for membranes templated on planar surfaces and on spherical surfaces. The transition releases free energy (exergonic) for membranes templated on cylindrical fibers of nanoscale radii. Merging of buds to obtain fewer buds of larger diameters results in a reduction in the elastic energy due to curvature of the population of buds. We show that a combination of budding and merging provides a net exergonic pathway for the assembly of GUVs on surfaces composed of nanoscale cylindrical fibers. We also show that the ratio of fiber radius to length that can support the exergonic formation of buds decreases with increasing adhesion potential. For lipid membranes on hydrophobic nanocellulose paper, this results in a switch from a net exergonic pathway to a net endergonic pathway.

The BNM model accounts for our observations that the yield of vesicles is highest on both the tracing paper and nanocellulose paper and for the dramatic reduction in yield on both these surfaces when the fibers are rendered hydrophobic. The BNM model is also consistent with observations of merging of vesicles buds on the surface. Since formation of larger vesicles requires the merging of many smaller buds, the model is consistent with the monotonically decreasing abundance of vesicles of larger diameters obtained from surface-assisted assembly techniques.

## Results and Discussion

### Surfaces used

The application of electric fields^29^, the swelling of the surfaces^32^, and the increased flux of water through permeable surfaces^36^ have been suggested to enhance the yields of GUVs. To control for these potential factors and to isolate the effect of surface curvature, we used seven different surfaces with varying properties. We used lab-made nanocellulose paper (NP), commercial artist-grade tracing paper (TP), silanized tracing paper (CH_3_-TP), regenerated cellulose dialysis membranes (RC), silanized glass slides (CH_3_-GS), pristine glass slides (GS), and indium tin oxide (ITO)-covered slides (electroformation, EF). The first five surfaces have not been used previously to assemble giant vesicles. The latter two surfaces are currently used to obtain giant vesicles^1^. Nanocellulose is obtained through chemical hydrolysis and high-pressure mechanical homogenization of regular cellulose fibers^37,38^. This treatment delaminates the cellulose fibers, which are tens of micrometers in diameter, into their constituent nanocellulose fibrils ~ 5 – 60 nm in diameter^37,38^. Dissolution and chemical regeneration of cellulose to form dialysis membranes result in smooth cellulose films devoid of fibrillar character^39^. The lab-made nanocellulose paper, commercial tracing paper, and regenerated cellulose dialysis membranes were insoluble in water, hydrophilic, permeable to water, and swell in water^37,38^. The glass and ITO-covered surfaces were impermeable and do not swell in water. Scanning electron microscopy (SEM) imaging showed that the nanocellulose paper, tracing paper, and silanized tracing paper were composed of entangled cylindrical nanofibers (Fig. 1a-c). The nanocellulose fibers were polydisperse in radius and length. The average radius of the fibers was *R =* 17 ± 6 nm, and the average length of the fibers was L= 5000 ± 2000 nm. The regenerated cellulose dialysis membrane (Fig. 1d), silanized glass slides, glass slides, and ITO-covered slides were planar and featureless. Grafting methyl groups onto the surfaces of the hydrophilic tracing paper and glass slides through silanization made the respective surfaces hydrophobic^40^. This surface treatment allowed us to probe the behavior of lipids on hydrophobic surfaces while preserving the geometry of the surfaces.

**Figure 1.**
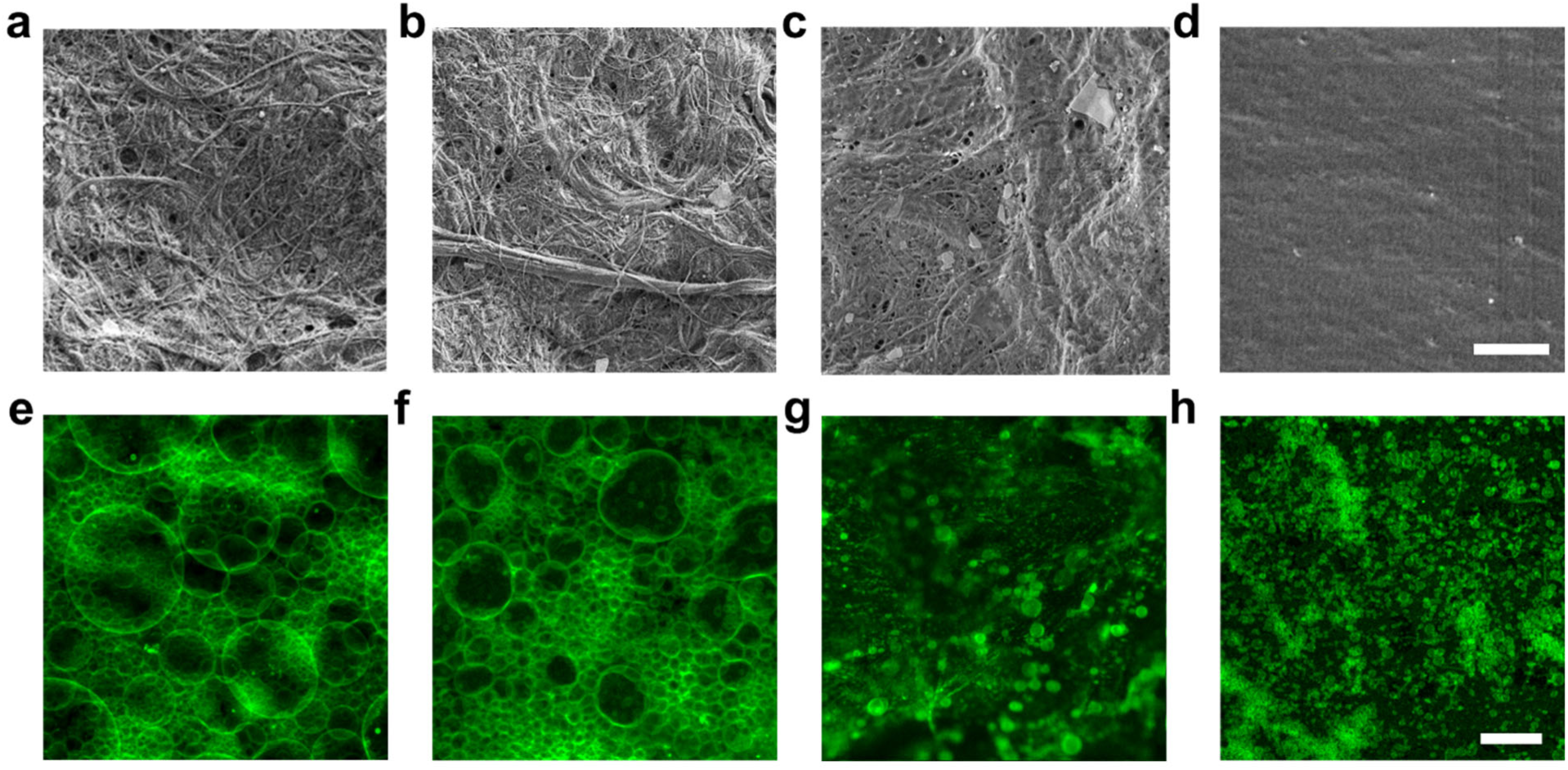
Characterization of the substrates and the assembled giant vesicle buds. (a-d) SEM images showing the microstructure of, (a) lab-made nanocellulose paper, (b) commercial tracing paper, **c** silanized tracing paper, and (d) regenerated cellulose dialysis membranes. (e-h) Confocal images of the templated lipid film after 2 hours in aqueous solutions on, **e** nanocellulose paper, (f) tracing paper, **(g)** silanized tracing paper, and (h) regenerated cellulose dialysis membranes. Scale bars: (a-d) 2 µm, (e-h) 50 µm.

We prepared thin films of lipids by drop-casting a solution of the zwitterionic phospholipid dioleoyl-*sn*-glycero-3-phosphocholine (DOPC) labeled with 0.5 mol % of the fluorescent sterol TopFluor-Cholesterol (TF-Chol) dissolved in chloroform. After removing trace solvents under vacuum, we incubated the surfaces in a 100 mM aqueous solution of sucrose. We applied an AC electric field to the ITO-covered glass slides to perform the electroformation technique. All the other surfaces were incubated without any further input of energy. Sample preparation for the papers and dialysis membrane was achieved easily using standard disposable 48-well plates, while for the glass slides and ITO-covered slides, custom chambers had to be assembled (See Supplementary Methods for further details). After 2 hours of incubation, we imaged the surfaces using high resolution confocal microscopy. Dense stratified layers of vesicle buds ranging in size from 1 – 150 µm covered the surfaces of the lab-made nanocellulose and commercial tracing papers (Fig. 1e,f). There were fewer buds on the surface of the silanized tracing paper (Fig. 1g) and regenerated cellulose dialysis membrane (Fig. 1h). We show similar images of the other surfaces in Supplementary Fig. 1.

### Hydrophilic surfaces with nanoscale cylindrical geometry produce high yields of GUVs

The local densities and sizes of buds varied widely even on a single surface (Fig. 2a, Supplementary Fig. 1). Thus, images of vesicular buds on the surface do not provide conclusive information on the effects of the different surfaces on yields. To obtain quantitative data, we harvested the vesicle buds from the surfaces. Harvesting allows statistical sampling since the buds from various locations on the surfaces become well-mixed in solution. We placed aliquots of the harvested vesicles in an imaging chamber filled with a 100 mM solution of glucose (Fig. 2b). The sucrose filled vesicles sediment to the bottom of the chamber since they have a higher density than the surrounding solution of glucose. After waiting for three hours, we obtained single-plane confocal tile scan images of the whole bottom surface of the imaging chamber. Confocal microscopy images have high spatial dynamic range. Each image allowed us to identify vesicles ranging from 1 µm in diameter up to hundreds of micrometers in diameter (Fig. 2c), i.e. the whole range of sizes classified as GUVs^1,2^. Using a custom image analysis routine, we obtain the counts, the distribution of sizes and the moments of the distribution of 𝒪(10^5^) − 𝒪(10^6^) vesicles per experiment (Fig. 2d). We performed five independent repeats for each surface.

**Figure 2.**
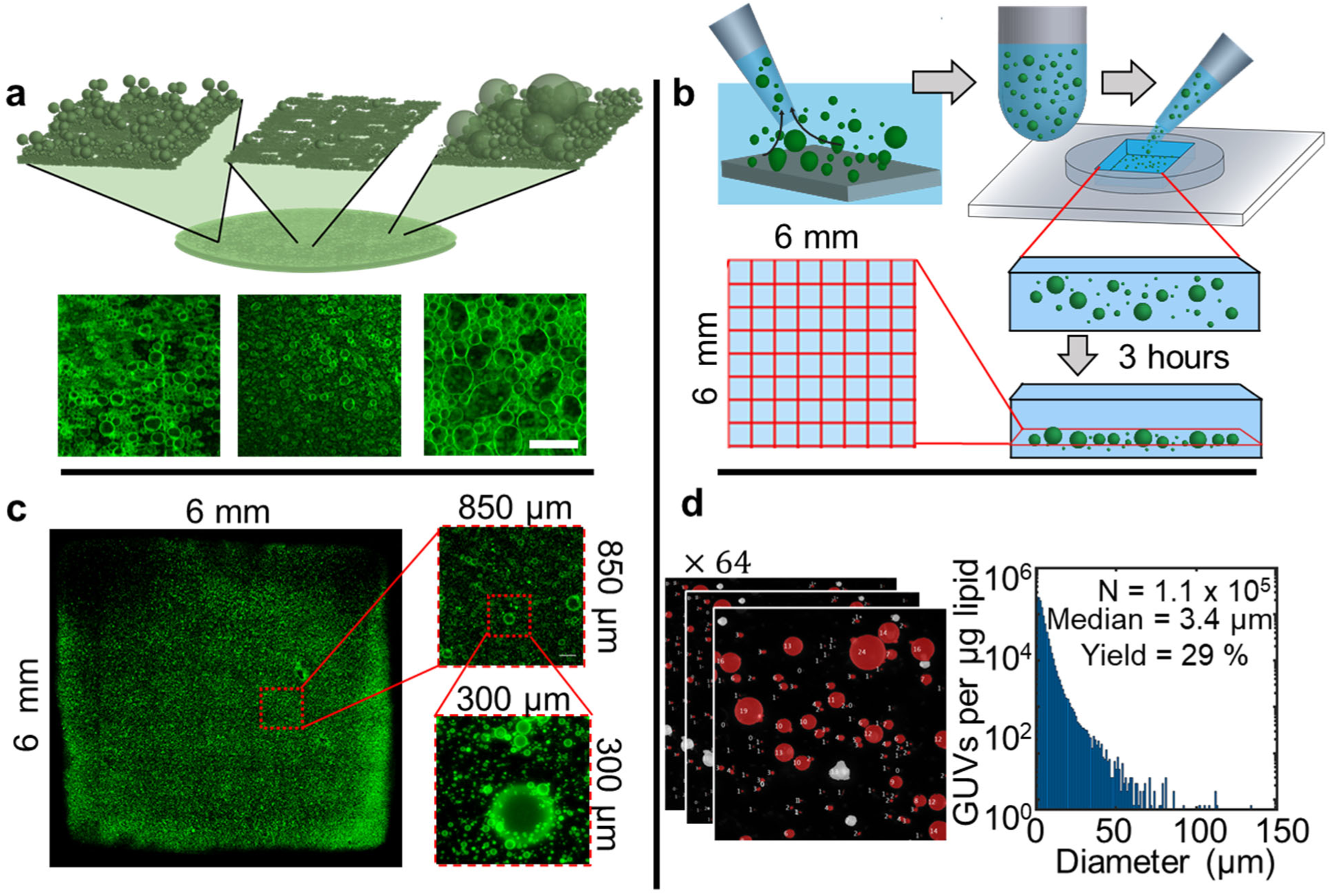
Quantitative analysis of GUVs (a) Cross-sectional schematic showing the large local differences in bud sizes and bud density on a substrate. The schematic was based on the experimental images in the bottom row, which are from assembly on a glass surface. Images from the other surfaces are shown in Supplementary Fig. 1. (b) Schematic showing the procedure for measuring the distribution of sizes and molar yield of GUVs. (Clockwise) Vesicle buds were harvested from the substrate and aliquots were introduced into a 6 mm × 6 mm square imaging chamber. After allowing the vesicles to sediment for 3 hours, we image the whole bottom surface of the chamber using single plane confocal microscopy. (c) Stitched confocal images of a typical tilescan of the vesicles in the chamber. The insets show progressive zoomed in views of a single tile and a region within the tile. Vesicles with diameters from 1 µm up to 150 µm are visible. (d) (Left) Example of processed images. The objects classified as GUVs are false colored red. The gray objects were not included in the count since they had intensities consistent with being multilamellar vesicles. An example of a histogram of GUV diameters obtained from a single repeat from tracing paper. The counts were normalized by the total mass of the DOPC deposited on the substrate. Bin widths are 1 µm. Note the logarithmic scale on the y-axis. Scale bar: 50 µm.

Fig. 2d shows a histogram of the diameters of the GUVs obtained from a sample of tracing paper. We show the histograms of the diameters of the GUVs obtained from the other surfaces in Supplementary Fig. 2, and report summary statistics in Supplementary Fig. 3. The distributions of diameters of the vesicles obtained from all the surfaces were right-skewed^41^, and showed monotonically decreasing counts as a function of increasing diameter — smaller vesicles were more abundant than larger vesicles (Supplementary Fig. 2a-g). Tracing paper and nanocellulose paper produced higher average counts of vesicles with larger diameters (Supplementary Fig. 3a) and produced populations of vesicles with larger median diameters (Supplementary Fig. 3b) and larger extreme diameters (Supplementary Fig. 3c) than the planar surfaces. The silanized surfaces produced lower counts of vesicles and vesicles with smaller diameters compared to their hydrophilic counterparts.

To allow comparison between surfaces, we define a ‘molar yield’, *Y*, of GUVs. The molar yield is the moles of lipids that compose the membranes of the GUVs normalized by the total moles of lipid deposited onto the surface. We obtain the molar yield from the confocal images using 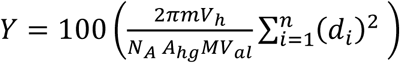. In this equation, *M* is the mass of lipid deposited on the surface, *m* is the molecular weight of the lipid, *V*_*al*_ is the volume of the aliquot in the imaging chamber, *V*_*h*_ is the volume of the harvested GUV suspension, is the number of GUVs in the imaging chamber, and *d*_*i*_ is the diameter of vesicle *i*.

Fig. 3a shows a stacked bar plot of the molar yields grouped by the surfaces with planar geometry (red) and surfaces composed of entangled nanocylinders (blue). The bars are divided by GUV size ranges, 1 *μm* ≤ *d* < 10 *μm* (small GUVs), 10 *μm* ≤ *d* < 50 *μm* (large GUVs), and *d* ≥ 50 *μm* (very large GUVs). The error bars are the standard deviation from the mean. We perform statistical tests on our data. An Anderson-Darling test for normality shows that the molar yield calculated from the independent repeats from all the surfaces were consistent with being drawn from a normal distribution (Supplementary Table 1). A Bartlett’s test showed that the samples had equal variances (Supplementary Table 1). Thus, the variation in the repeats is consistent with the additive effects of multiple independent processes and is unlikely to be due to systematic effects^42^. To assess statistical significance of the effect of the surfaces on molar yields, we perform a balanced one-way analysis of variance (ANOVA). The ANOVA showed that at least one of the surfaces had a significant effect on the yield of GUVs [*F*(6,28) = 90.53, *p =* 5.06 × 10^−17^]. We performed Tukey’s Honestly Significant Difference (HSD) post hoc tests to determine statistical significance between pairs of surfaces. A p-value of < 0.05 was considered significant. See Supplementary Table 2 for the ANOVA table and the results of the Tukey’s HSD.

**Figure 3.**
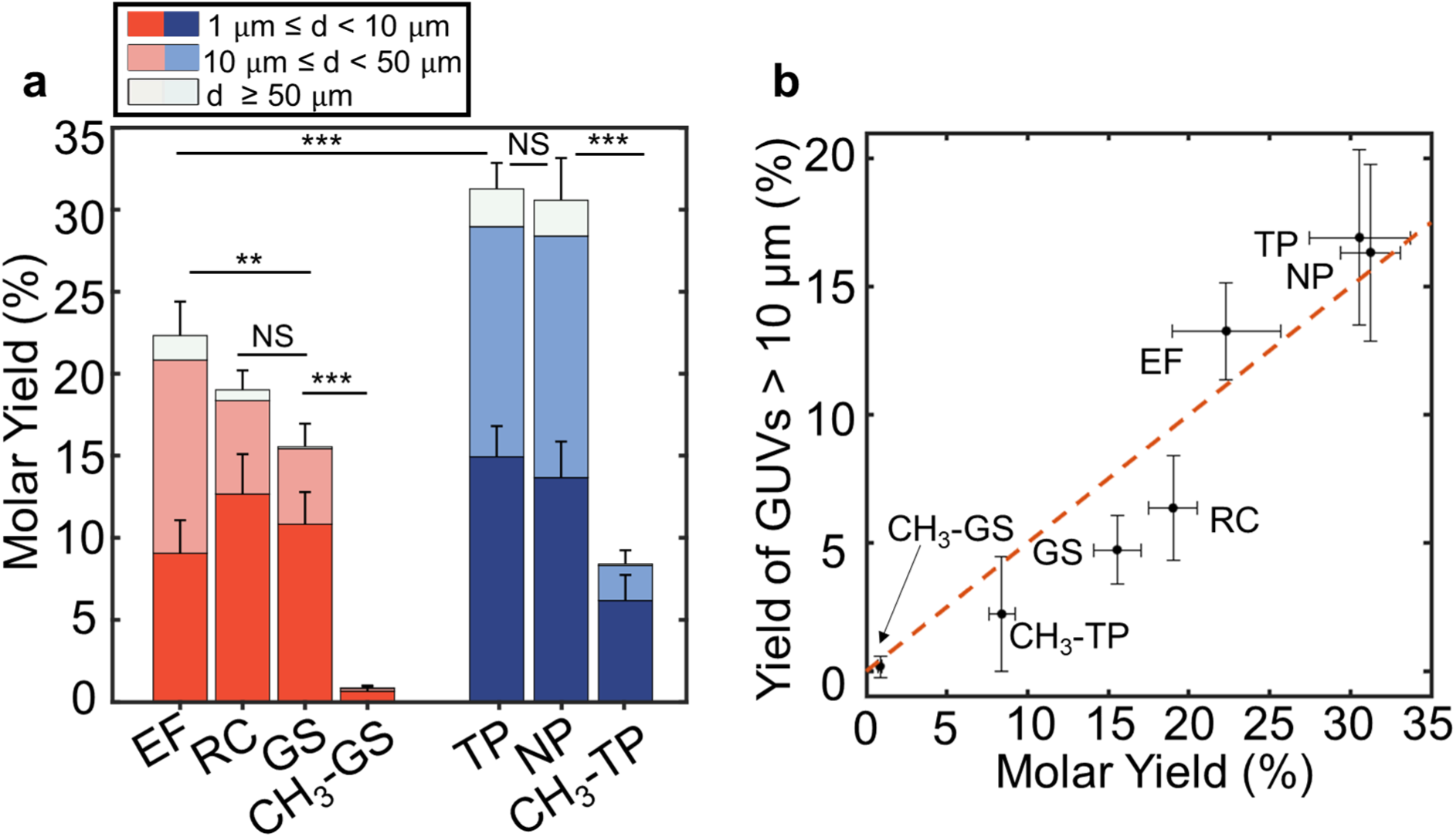
Molar yields of GUVs obtained from planar substrates and substrates composed of cylindrical nanofibers. (a) Stacked bar plots of the molar yields of GUVs obtained from growth on planar substrates (red) and substrates composed of cylindrical nanofibers (blue). Each bar is an average of *N* = 5 independent repeats and the error bars are the standard deviation from the mean. The planar substrates are electroformation on ITO-covered slides (EF), regenerated cellulose dialysis membranes (RC), glass slides (GS), and silanized glass slides (CH_3_-GS). The substrates composed of cylindrical nanofibers are tracing paper (TP), lab-made nanopaper (NP), and silanized tracing paper (CH_3_-TP). Each bar is split into three regions corresponding to the fraction of vesicles with diameter ranges as described in the legend. Statistical significance was determined using a one-way ANOVA and Tukey’s HSD post-hoc tests. *\* = p* < 0.05, *\*\* = p* < 0.01, *\*\*\* = p* < 0.001, NS = not significant. (b) Scatter plot showing the yield of GUVs with diameters larger than 10 µm versus the total molar yield of GUVs. The orange dashed line corresponds to half the GUVs being larger than 10 µm. The *x-* and *y-* error bars are the standard deviation from the mean.

The molar yield of GUVs from nanocellulose paper and tracing paper both ranged between 26% to 36% with a mean of 31%. The molar yield of GUVs from both surfaces were significantly higher than the molar yield of GUVs obtained from the glass slides, regenerated cellulose dialysis membranes, and ITO-covered glass slides (electroformation) which were 16 ± 1%, 19 ± 2%, and 22 ± 2% respectively (all p < 0.001, ***). The molar yield of GUVs obtained from the regenerated cellulose dialysis membranes was not significantly different than glass (p=0.383). The approximately 6% higher molar yield of GUVs obtained through electroformation compared to glass was statistically significant, albeit with a lower confidence level of p = 0.005, **. The difference in yield between electroformation and regenerated cellulose dialysis membranes was not statistically significant (p = 0.439). The silanized glass slide and silanized tracing paper surfaces had mean molar yields of 1 ± 0.4% and 8 ± 2% respectively. This approximately 16-fold reduction in yield for silanized glass and ~ 4-fold reduction in yield for silanized tracing paper compared to their respective hydrophilic surfaces was highly significant (both p < 0.001, ***).

Summarizing these observations, we conclude that, i) surfaces consisting of nanoscale cylinders produce significantly higher yields of GUVs compared to planar surfaces, ii) the increased permeability of water through a surface does not lead to a statistically significant increase in the molar yield of GUVs compared to an impermeable surface, iii) the application of an electric field leads to a statistically significant increase in the yield of GUVs compared to gentle hydration on glass slides, and iv) hydrophobic modification of surfaces results in a significant reduction in GUV yields.

Fig. 3b shows a scatter plot of the mean yield of GUVs with diameters greater than 10 µm (large GUVs) versus the mean yield of the population. The vertical and horizontal error bars are the standard deviation from the mean. The dashed orange line shows the values at which half of the vesicles in a population are classified as large. The yield of large vesicles is highly correlated with the total yield of vesicles (Pearson’s R= 0.9595, p=6.194 × 10^−4^). For surfaces that had low yields, less than half of the vesicles were larger than 10 µm. For surfaces that had high yields, more than half of the vesicles were larger than 10 µm. Thus, the fraction of large vesicles in a population is proportional to the total yield of vesicles in the population. We surmise that obtaining populations consisting exclusively of large vesicles from surface-assisted hydration is highly unlikely.

### The budding and merging model explains the effects of surface geometry and surface hydrophobicity

Current proposed mechanisms^29,31,43–45^ for the formation of GUVs cannot explain our data. Based on our observations, we propose that GUVs form on lipid films templated on surfaces through the process of bud formation followed by the merging of the buds (budding and merging, BNM).

To determine the energy for bud formation, we model the free energy of a vesiculating membrane that is templated on a surface using Equation (1):

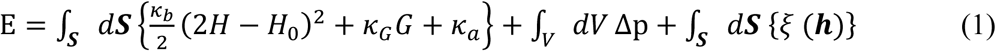

In this equation, *κ*_*b*_ is the bending modulus, is the Gaussian bending modulus, 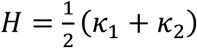 is the mean curvature where *κ*_1_ and *κ*_2_ are the principal curvatures on the surface, *G* = *κ*_1_ *κ*_2_ is the Gaussian curvature, *H*_0_ is the spontaneous curvature, *κ*_*a*_ is the area expansion modulus, Δp is the osmotic pressure difference, and *ξ*(***h***) is the microscopic interaction potential normal to the surface of the membrane. The microscopic interaction potential can be traced to intermolecular electrostatic, van der Waals, and structural forces^46^. The magnitude of these forces depends on the distance, ***h***, between the membrane and the surface^46^.

The first integral on the right-hand side is the elastic energy of the membrane using the Helfrich harmonic approximation^47–49^. The expression penalizes bending from *H*_0_ = 0 of a symmetric bilayer membrane and for any stretching of the membrane. The Gaussian curvature is invariant absent a topological change^50^. Since the buds remain attached to the surface, the Gaussian curvature of the membrane is invariant during the process of budding (See Supplementary Fig. 4 and Supplementary Text for a discussion of the morphology and connectivity of the buds). A region of negative Gaussian curvature develops at the neck of the bud that equals the positive Gaussian curvature of the spherical bud^50^. We thus only consider changes in elastic energy of the membrane due to changes in the mean curvature. The second term in Equation (1) accounts for pressure-volume work. Osmotic pressure variations can arise from differences in the distribution of lipid counterions^27^. Since all the surfaces were templated identically with DOPC, we take Δ*p* ≈ 0. The third term in Equation (1) accounts for the interaction between the membrane and the surface. In addition to the bare surface, membranes organized as multilayers could have a bilayer, a depleted bilayer^51^, or a monolayer as a ‘surface’. Following^48,49^, we ignore microscopic details and replace the microscopic interaction potential, *ξ*(***h***) with an effective adhesion contact potential, *ξ*. The adhesion potential varies with the identity of the surface and can range over five orders of magnitude^46,51^. We show characteristic values for these parameters for phosphocholine lipid bilayers in Supplementary Table 3.

The energy change, ∆*E* = *E*_2_ − *E*_1_ for the membrane templated on the surface to transition to a spherical bud at a constant surface area, ***S***_**1**_ = ***S***_**2**_ is given by Equation (2):

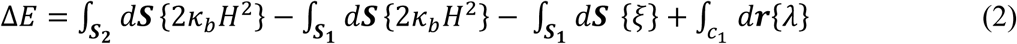

The last term in Equation (2) introduces a constraint for a section of the membrane to transition into a spherical bud at a constant area. If there is a lipid source, the membrane can transition to a spherical bud without requiring breaks by recruiting lipids from the source (Fig. 4a,c). In the absence of a lipid source, the membrane must form breaks, with an edge energy *λ* to allow the lipids to reconfigure to form the spherical bud (Fig. 4b,d).

**Figure 4.**
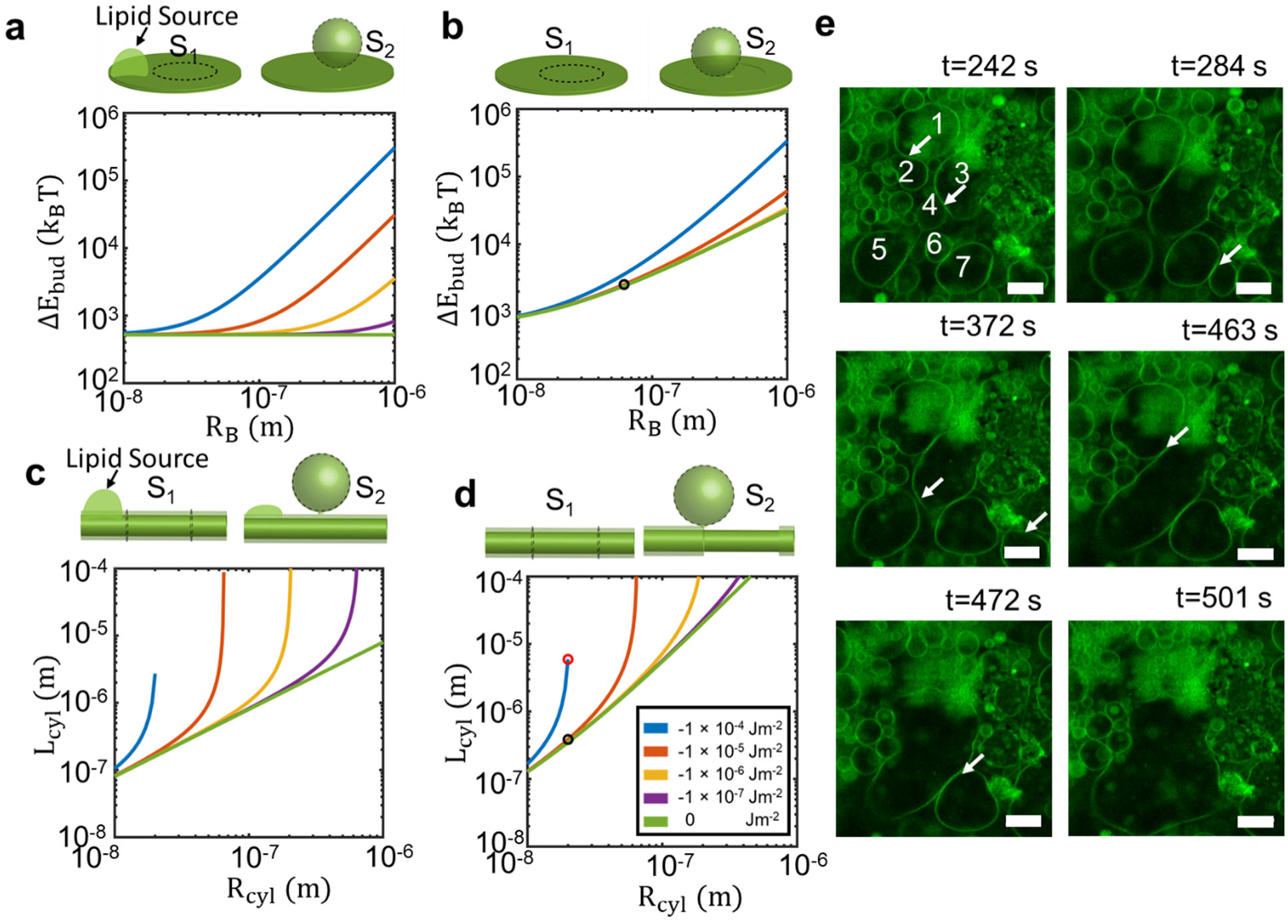
The budding and merging model. (a-d) The schematics show the configuration of the membrane templated on the surface transitioning to a spherical bud at a constant surface area, ***S***_**1**_ = ***S***_**2**_. Plots of the change in free energy as a function of bud size when the membrane transitions from a plane to a spherical bud when, (a) there is a lipid source, (b) when there is no lipid source. All energies are relative to the thermal energy scale, *k*_*B*_*T*, and are significantly above zero. Plots of isoenergy (Δ*E*_*cyl*_ = 0) lines in *R*_*cyl*_ − *L*_*cyl*_ phase space when a membrane transitions to a spherical bud, (c) when there is a lipid source, (d) when there is no lipid source. The red and black circles in (d) represent the test radius of *R*_*cyl*_ = 20 nm and length *L*_*cyl*_ = 2000 as described in the main text. The colors of the line denote different adhesion energies as described in the legend in (d). Note the logarithmic scale for both the *x-* and *y-* axes of these plots. **e** Stills from a confocal time-lapse of 7 vesicle buds merging over the course of ~ 5 minutes. The first image was captured 4 minutes after initial immersion of the lipid-coated paper in water. The white arrows indicate vesicle walls that merge. Scale bars 10 µm.

### The free energy change due to budding can be negative on nanoscale cylinders

The curvature elastic energy of a spherical bud is 8*πκ*_*B*_ and is independent of the size of the bud^47^. The change in energy for lifting a planar disk, with a radius of *R*_*disk*_ off of the surface to form a bud of radius 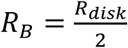 is given by Equation (3):

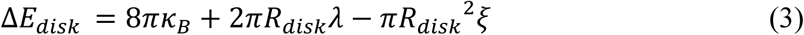

Fig. 4a shows this change in energy as a function of bud size for transitions with a lipid source for values of adhesion energies *ξ* = 0, −1 × 10^−7^, −1 × 10^-6^, −1 × 10^−5^, and −1 × 10^−4^ J m^−2^. Fig. 4b shows the change in energy as a function of bud size for transitions without a lipid source. Note the logarithmic scale on both the *x* and *y* axes. The change in free energy is positive (endergonic) for all bud sizes and is significantly above the thermal energy scale *κ*_*B*_*T* = 4.11 × 10^−21^ J at a temperature, T=298 K. The Boltzmann constant is *κ*_*B*_. Since adhesive interactions and the edge energy scales as 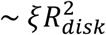 and 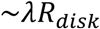, smaller buds have a lower energy of formation.

For a cylindrical section of membrane on a cylindrical fiber, there are multiple combinations of radii,*R*_*cyl*_ and length, *L*_cyl_ of the membrane that can produce a spherical bud with radius, 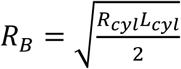. The change in energy for forming a spherical bud is given by Equation (4):

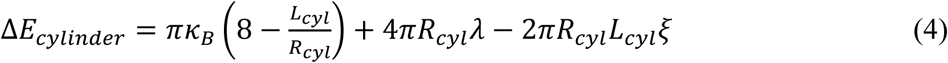

For a given *R*_*B*_, longer *L*_*cyl*_ and smaller *R*_*cyl*_ result in a larger change in energy. This is because the elastic potential energy due to curvature of the membrane scales as 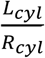. The adhesive interactions and the edge energy scales as ~2*πR*_*cyl*_*L*_*cyl*_*ξ* and ~4*πR*_*cyl*_*λ*. Intriguingly, bud formation on nanocylinders of different lengths can result in zero and even negative change in free energies (exergonic) (Fig. 4c). The colored lines are isoenergy lines where Δ*E*_*cylinder*_ = 0 plotted in *R*_*cyl*_ − *L*_*cyl*_ phase space. These lines delineate the endergonic and exergonic regions for a given adhesion potential (Fig. 4c). The magnitude and sign of Δ*E*_*cylinder*_ at each coordinate can be obtained by substituting, (*R*_*cyl*_,*L*_*cyl*_,*ξ*) into Equation (4). The bud size that is formed at that coordinate is given by 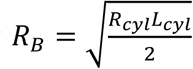. Values to the left and above of the isoenergy line for a given *ξ* have exergonic changes in free energies. Values to the right and below the isolines result in increasingly endergonic changes in free energies.

The model shows that attractive adhesion potentials of increasing magnitude results in a smaller range of fiber radii that can support the exergonic growth of buds. Formation of buds with breaks in the membrane decreases the size of the exergonic region (Fig. 4d). Interestingly, we find that the exergonic formation of spherical buds is peculiar to fibers with cylindrical geometry. Membranes templated on spherical particles, despite their curvature, have an overall positive change in free energy (See Supplementary Text).

### Assembly of giant vesicle buds proceeds through merging of buds on the surface

We estimate the number of buds that form per unit area of surface, 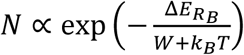, where, 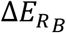 is the free energy change of a bud of size *R*_*B*_ and *W* is the external energy available in the system. The process of hydration introduces external energy from hydrodynamic flows or temperature gradients^43^. These sources of energy have been proposed to cause the formation of giant vesicles^43^. Due to their lower free energy of formation, nanoscale buds should be exponentially more abundant than micrometer scale buds on the surfaces. Note that unlike on planar surfaces, thermal energy can drive the formation of buds on surfaces with nanoscale cylindrical geometry since the change in free energy is negative (Fig. 4c,d).

If nanoscale buds are exponentially favored, why do we obtain buds of sizes that range up to hundreds of micrometers (Fig. 1)? External sources of energy could cause the formation of large buds^43^. However, since inputs of external energy is expected to be random, input of external energy alone cannot explain the highly statistically significant increase in yields of vesicles obtained from surfaces with nanoscale cylindrical geometry (Fig. 3a,b).

To obtain insight, we observe the dynamics of buds on the surfaces using time-lapse confocal microscopy. We find that vesicular buds with diameters greater than 1 µm and as large as 10 µm were present within 4 minutes of hydrating the dry lipid-coated paper (Fig. 4e). Further, the buds evolved by merging on the surface. Smaller buds that were initially tens of micrometers apart form a single larger bud through a series of cascading merging events over the course of 5 minutes (Fig. 4e). Merging results in the formation of buds that span many nanofibers (compare the scales in Fig. 1a,b to Fig. 1d,e). We note that the total curvature energy of the vesiculating membrane is only dependent on the number of spherical buds and not on the size of the buds. Thus, merging while maintaining a constant bud area to form fewer buds of larger sizes results in the reduction of the free energy of the vesiculating membrane.

Since we cannot directly observe the nanoscale buds with confocal microscopy, we posit that the process of merging observed for the micron-scale buds extends to the nanoscale. This supposition gives us the driving force for the formation of large micrometer scale buds from nanoscale buds. We consider an idealized pathway for forming a GUV bud 10 µm in radius on a surface consisting of nanocylindrical fibers with *R*_*cyl*_ = 20 nm and *L*_*cyl*_ = 2000 nm. We use an adhesion energy of −1 × 10^−5^ J m^−2^ for DOPC bilayers interacting with each other^46^. We consider the case where there is no lipid source and thus transient breaks in the membrane must occur during budding. We obtain a characteristic length *L*^∗^ ≈ 384 nm where the budding energy Δ*E*_*cylinder*_ = 0 (red circle in Fig. 4d). The membrane on each fiber can form ~ 5 buds with a characteristic bud radius 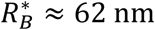. 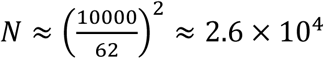 of these nanobuds must merge to obtain a single bud 10 µm in radius. Nominally the system releases 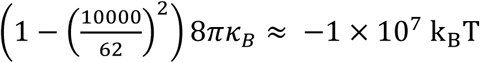 through budding and merging. The highly exergonic nature of this idealized process suggests that nanobuds are short lived on the surfaces. This result is consistent with the rapid emergence of micrometer-scale buds on the surface of the paper (Fig. 4e). Note that viscous dissipation and other barriers will reduce the net energy change. In contrast, since the formation of buds on planar surfaces is always endergonic, there is no thermally favored characteristic bud size. Using 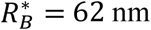 as a comparison, the budding energy on a plane is ~ 2533 k_B_ T per bud. The net energy change for the formation of a vesicle through budding and merging on a planar surface is ≈ 8 × 10^7^ k_B_ T.

An increase in the adhesion potential by an order of magnitude to −1 × 10^−4^ J m^−2^ gives *L*^*^ = 5920 nm and 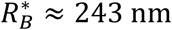 on the nanofiber surface (black circle, Fig 4d). This value of *L*^∗^ is ~ 3 times larger than the L=2000 nm of the idealized nanofibers that we chose. Thus, despite the nanoscale curvature of the fibers, formation of buds on this surface becomes endergonic. This large effect of the magnitude of the adhesion potential on the energetics of bud formation is consistent with our observation of the significant effect that silanization has on vesicle yields both on surfaces with planar geometry and on surfaces consisting of nanoscale fibers (Fig. 3a). This is because the adhesion potential for membranes on hydrophobic surfaces can be as high as −1 × 10^−1^ J m^−2^ due to the formation of depleted membranes that expose hydrophobic regions^51^. On a surface consisting of entangled fibers of various lengths and radii, the effect of locally different adhesion potentials and different dimensions of fibers could be reflected by having buds forming only on a fraction of the nanofibers.

We note that the budding and merging model is consistent with the higher yields obtained through electroformation compared to gentle hydration on glass (Fig. 3a). The electric field inputs energy into the system which can drive bud formation. Such a source of energy is absent for gentle hydration on glass and regenerated cellulose dialysis membranes. Further, electric fields reduce the adhesion between phosphocholine bilayers^29^, which in our model results in the formation of a larger number of buds for a given amount of external energy. Clearly however, nanoscale cylindrical geometry is more effective in increasing vesicle yields than the application of electric fields to lipid films.

Merging as the predominant mechanism of bud growth is consistent with the high positive correlation between the fraction of large vesicles and the total amount of vesicles (Fig. 3b). It is rational that the formation of larger numbers of nanobuds at a higher density on the surface will lead to a greater number of merging events. Increased merging results in the formation of larger vesicles on the surface. Further, many merging events must occur to obtain vesicles of larger diameters. This scenario is consistent with our observations that very large vesicles are statistically rare (Supplementary Fig. 2), while smaller vesicles are more abundant.

### Nanopaper has practical methodological and cost advantages for small and larger scale assembly of GUVs

Our data allows consideration of practical aspects of surface-assisted assembly of GUVs. We calculate the cost of substrates per vesicle (Fig. 5a) using the lowest posted prices from the websites of large multinational suppliers of scientific materials (Supplementary Table 4). The continuous lines are the mean substrate cost for a given vesicle size and the shaded region around the lines are the standard deviation from the mean. Obtaining GUVs using tracing paper has the lowest cost per vesicle of all sizes. The cost reduction is stark, particularly for vesicles of larger diameters since they occur at a lower abundance. For example, producing a single 100 µm diameter GUV using electroformation costs USD 0.16. The cost is more than 100,000× lower when tracing paper (USD 0.0000013) is used as a surface. Note that for gentle hydration on glass, vesicle sizes above 60 µm were not accessible. We use a prototypical example of producing 1 liter of artificial blood, a still unsolved challenge^52,53^, to relate this cost analysis to the required scales for potential synthetic cell or artificial tissue applications. The typical concentration of erythrocytes in a healthy adult male^54^ is 4.92 × 10^12^ cells/L. The biconcave cells have an equivalent mean spherical diameter of 5.6 µm^54^. Obtaining 1 L of GUVs with diameters between 5.0 and 5.9 µm at a concentration of 4.92 × 10^12^ GUVs/L using electroformation would require a surface area of 530 m^2^. The cost of the surface is approximately USD 12,000,000. In contrast, obtaining the same number of GUVs using tracing paper would require a surface area of 150 m^2^ at a cost of USD 120.

**Figure 5.**
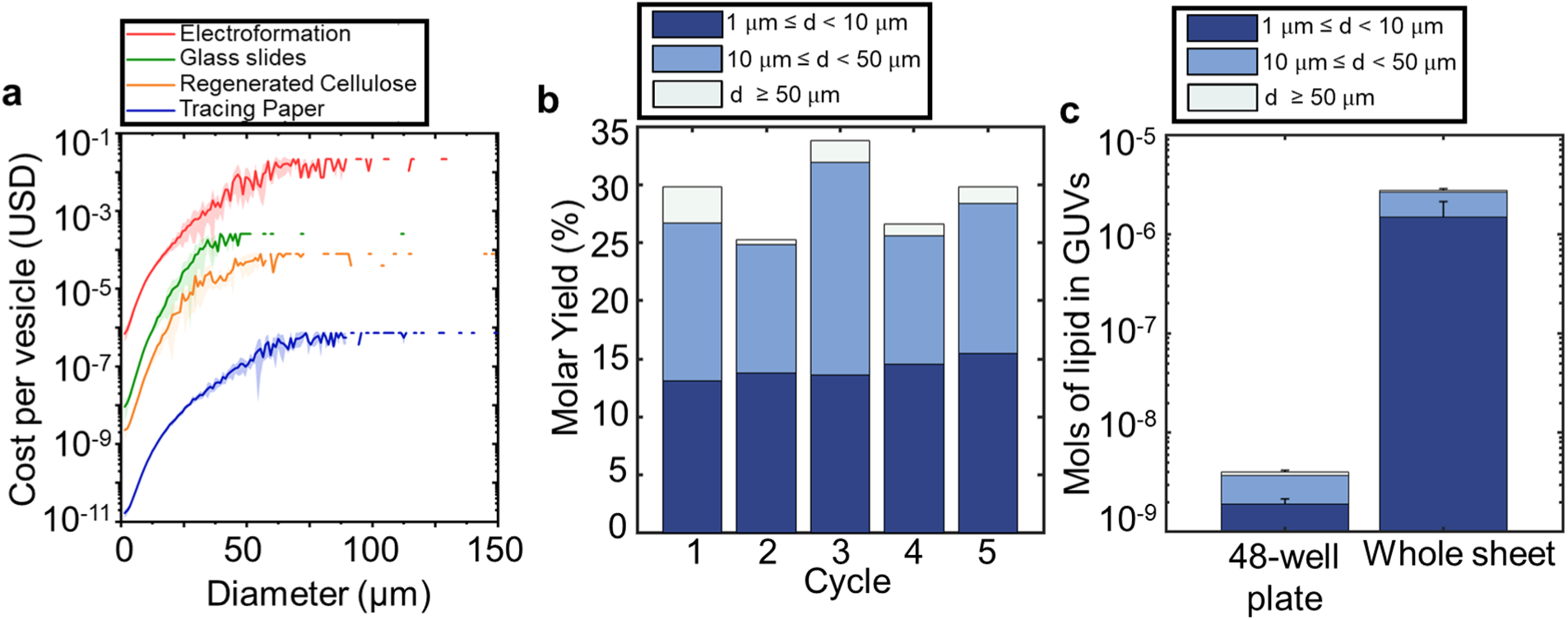
Analysis of costs and scalability. (a) Line plots showing the average cost of the substrate in US dollars to produce a single vesicle of a given diameter using electroformation (red), glass slides (green), regenerated cellulose dialysis membranes (orange), and tracing paper (blue). Note the logarithmic scale on the y-axis. The line shows the mean of *N* = 5 independent repeats and the shaded region around the lines are the standard deviation from the mean. Bin widths are 1 µm. Electroformation had the highest costs per vesicle and tracing paper had the lowest costs per vesicle. (b) Stacked bar plot of the molar yield of GUVs harvested 5 times cyclically by reusing a single piece of tracing paper. (c) Stacked bar plot of the mols of lipid harvested as GUVs using a 48-well plate experiment and a larger-scale whole sheet experiment. Note the logarithmic scale on the *y-*axis which compresses the stacks within the bars.

In addition to being the cheapest single-use surface, the high tensile strength of nanocellulose^37,38^ makes tracing paper resilient to mechanical insults. This resiliency allows sterilization and rigorous cleaning before use or reuse. Fig. 5b shows stacked bar plots of the molar yields of GUVs obtained from five cycles of use of a single piece of tracing paper. The molar yields are unchanged within experimental variability in each of the five cycles. SEM images of the tracing paper after the fifth cycle of use were indistinguishable from the paper after the first use (Supplementary Fig. 5). Thus, the tracing paper could be reused many more times, further lowering fixed substrate costs. For example, reusing the paper five times reduces the amount of surface needed for making 4.92 × 10^12^ GUVs with diameters between 5.0 and 5.9 µm to 30 m^2^ and the cost to USD 24.

The manipulability of paper allows simple scale-up. We scale our typical process which was optimized for 48-well plates by using a whole sheet of tracing paper (12-inch × 9-inch), a commercial air-brush suitable for spraying harsh volatile solvents, and a 13-inch × 9-inch baking tray as a fluid receptacle. Fig. 5c shows a stacked bar plot of the total number of GUVs obtained from the scaled-up experiment compared to the 48-well plate experiment. Note the logarithmic scale on the *y*-axis which compresses the stacks within the bars. We obtain about 600 times more GUVs, measured as the mols of lipids harvested, through the larger format experiment.

## Conclusions

The results presented here have both practical and fundamental implications. The budding and merging model accounts for the effect of surface curvature and hydrophobicity on the yield of vesicles and is consistent with the right-skewed distribution of vesicle diameters in a population. Our introduction of the concept of a molar yield of GUVs allows statistically rigorous comparisons between different surfaces. Measurements of the molar yield can be extended to studies of other parameters that might affect the formation of GUVs such as the temperature or the type of lipid. Practically, the low cost of paper and the ability to use standard laboratory plasticware, such as multiwell plates and Eppendorf tubes, makes surface-assisted assembly using nanocellulose paper highly accessible in the research laboratory. The wide availability of industrial machines that print solutions of volatile solvents and that handle paper on a massive scale^39^ suggests easy adaptation to current industrial manufacturing practices. Our results thus address practical barriers that currently impede the promising use of GUVs as vehicles for the delivery of drugs, the manufacturing of synthetic cells, and the assembly of artificial tissues at scale.

## ASSOCIATED CONTENT

### Supporting Information

The Supporting Information is available free of charge on the ACS Publications website at DOI:

Supplementary text describing the morphology of the buds on the surfaces and mathematical details of the method. Supplementary methods. Figures showing: Confocal images of buds on the different surfaces. Histograms of GUVs from different surfaces. GUV counts, median diameters, and extreme diameters. FRAP experiments showing connectivity of buds on the surface. SEM images showing the microstructure or reused tracing paper. Table of p-values for statistical tests. Table of physicochemical properties of phosphocholine. Table of costs of substrates. Figure S1-S5, Table S1-S4. (PDF).

## AUTHOR INFORMATION

### Author Contributions

The manuscript was written through contributions of all authors. All authors have given approval to the final version of the manuscript.

## Acknowledgements

This work was funded by the National Science Foundation through NSF CAREER DMR-1848573, NSF CBET-1512686, and NSF-CREST: Center for Cellular and Biomolecular Machines at the University of California, Merced (NSF-HRD-1547848). The data in this work was collected, in part, with a confocal microscope acquired through the National Science Foundation MRI Award Number DMR-1625733, and a scanning electron microscope acquired through NASA Grant NNX15AQ01A.

