## Supplementary Information for "Nanoscale curvature promotes high yield spontaneous formation of cell-mimetic giant vesicles"

### Supplementary Text

#### 1. Morphology and connectivity of the vesicle buds on the surfaces

Vesicles are topologically isolated membranes (Supplementary Fig. 4a). In contrast, vesicle buds are continuous membrane protuberances that are connected to the lipid film on the substrate and to each other (Supplementary Fig. 4b). We use fluorescence recovery after photobleaching (FRAP) to distinguish isolated vesicles from connected vesicular buds. The distinction between buds and topologically isolated vesicles is critical for understanding the mechanism of formation of GUVs. We hypothesized that isolated vesicles will be permanently photobleached in a FRAP experiment since the closed membranes are not connected to sources of unbleached TF-Chol (Supplementary Fig. 4a). Photobleached vesicular buds can recover their fluorescence intensity since the membranes are connected to sources of unbleached TF-Chol (Supplementary Fig. 4b). Supplementary Fig. 4c and d shows selected stills and the recovery curves of FRAP experiments performed on a close-packed layer of isolated vesicles (Supplementary Fig. 4c) and on a close-packed layer of vesicle buds on the surface of tracing paper (Supplementary Fig. 4d). Prior to bleaching, both the vesicles and the vesicle buds appear similar in confocal images. Upon bleaching, the fluorescence intensity of the isolated vesicles did not recover (Supplementary Fig. 4c). The fluorescence intensity of the buds on the surface of the tracing paper recovered (Supplementary Fig 4d). This result proves that the morphology of the film on the surface is consistent with a continuous layer of connected vesicular buds.

### 2. Calculations of the change in energy for planar, cylindrical, and spherical substrate geometries

#### 2.1 Change in energy of a disk with radius $R_{disk}$ to a bud with $R_{bud}$

$$E_1 = \pi {R_{disk}}^2 \xi \tag{2.1.1}$$

$$E_2 = 8\pi\kappa_B + 2\pi R_{disk}\lambda \tag{2.1.2}$$

$$\Delta E_{disk} = E_2 - E_1 = 8\pi\kappa_B + 2\pi R_{disk}\lambda - \pi {R_{disk}}^2 \xi \tag{2.1.3}$$

Setting $\Delta E = 0$ and solving for $R^*_{disk}$ gives us:

$$R^*_{disk} = \frac{\lambda + \sqrt{(\lambda^2 + \xi + 8\kappa_B\xi)}}{\xi} \tag{2.1.4}$$

This value is always negative for attractive interactions. Thus, there are no physical values of $R_{disk}$ where $\Delta E \leq 0$ for $\xi \leq 0$, e.g. for attractive or zero adhesion potentials. If there are lipid sources, no breaks in the membrane are required. In this case, $\lambda = 0$.

Area constraint:

$$R_{disk}{}^2 = 2R_B^2 \tag{2.1.5}$$

**2.2 Change in energy from a cylinder of radius $R_{cyl}$ and length $L_{cyl}$ to a sphere of radius $R_B$**

$$E_1 = \frac{\pi\kappa_B L_{cyl}}{R_{cyl}} + 2\pi R_{cyl} L_{cyl}\xi \tag{2.2.1}$$

$$E_2 = 8\pi\kappa_B + 4\pi R_{cyl}\lambda \tag{2.2.2}$$

$$\Delta E_{cylinder} = E_2 - E_! = \pi\kappa_B\left(8 - \frac{L_{cyl}}{R_{cyl}}\right) + 4\pi R_{cyl}\lambda - 2\pi R_{cyl} L_{cyl}\xi \tag{2.2.3}$$

Setting $\Delta E = 0$ and solving for $L^*$ gives:

$$L^* = \frac{8\lambda R^2 + 8\kappa_B R}{\kappa_B + 2\xi\, R^2} \tag{2.2.4}$$

For attractive interactions, i.e. when $\xi$ is negative, $L*$ is undefined when $\kappa_B = 2\xi\, R^2$ and becomes negative when $|\kappa_B| < |2\xi\, R^2|$ which are unphysical solutions. If there are lipid sources, no breaks in the membrane are required. In this case, $\lambda = 0$.

Area constraint:

$$R_{cyl} L_{cyl} = 2R_B^2 \tag{2.2.5}$$

**2.3 Change in energy from a hemispherical membrane on a bead of radius $R_{bead}$ to a spherical bud**

$$E_1 = 4\pi\kappa_B + 2\pi R_{bead}^2\, \xi \tag{2.3.1}$$

$$E_2 = 8\pi\kappa_B + 4\pi R_{bead}\lambda \tag{2.3.2}$$

$$\Delta E_{sphere} = 4\pi\kappa_B + 4\pi R_{bead}\lambda - 2\pi R_{bead}^2\, \xi \tag{2.3.3}$$

$$R^*_{bead} = \frac{\lambda - \sqrt{(\lambda^2 + 2\kappa_B \xi)}}{\xi} \quad (2.3.4)$$

This value is always negative for attractive interactions. Thus, there are no value of $R_{bead} > 0$ where $\Delta E < 0$ for $\xi \leq 0$, e.g. for attractive or zero adhesion potentials. If there are lipid sources, no breaks in the membrane are required. In this case, $\lambda = 0$.

Area constraint

$$R^2_{bead} = 2R^2_B \quad (2.3.5)$$

### 3. Supplementary Methods

#### 3.1. Materials and Chemicals

**Materials:** We purchased indium tin oxide (ITO) coated-glass slides (25 × 25 mm squares, surface resistivity of 8-12 Ω/sq) from Sigma-Aldrich (St. Louis, MO). We purchased a 3 wt % aqueous slurry of nanofibrillated cellulose from the University of Maine Process Development Center. We purchased artist grade tracing paper (Jack Richeson & Co., Inc.), circular hole punches (EK Tools Circle Punch, 3/8 in.), square hollow punch cutters (Amon Tech), a Paasche Gravity Feed Airbrush Kit (Model TG-3W) and Paasche Compressor system (Model D3000R) from Amazon Inc. (Seattle, WA). We purchased Fisherbrand Regenerated Cellulose Dialysis Tubing (MWCO-12,000-14,000), glass coverslips (Gold Seal™ , 22 mm x 22 mm), premium plain glass microscope slides (75 mm x 25 mm), and glass Petri dishes (Pyrex™, 150 mm diameter**)** from Thermo Fischer Scientific (Waltham, MA).

**Chemicals:** We purchased sucrose (BioXtra grade, purity ≥99.5%), glucose (BioXtra grade, purity ≥99.5%), Triton X-100, and casein from bovine milk (BioReagent grade), toluene (anhydrous, 99.8%) from Sigma-Aldrich (St. Louis, MO). We purchased chloroform (ACS grade, purity ≥99.8%, with 0.75% ethanol as preservative), acetone (ACS grade, purity ≥99.5%), methyltrichlorosilane (ACS grade, purity ≥99.5%) from Thermo Fischer Scientific (Waltham, MA). We obtained 18.2 MΩ ultrapure water from an ELGA Pure-lab Ultra water purification system (Woodridge, IL). We purchased 1,2-dioleoyl-sn-glycero-3-phosphocholine (18:1 (Δ9-cis) PC (DOPC)), 23-(dipyrrometheneboron difluoride)-24-norcholesterol (TopFluor-Cholesterol), from Avanti Polar Lipids, Inc. (Alabaster, AL).

**Fabrication of nanocellulose paper:** We fabricated nanocellulose paper in the laboratory using solution casting[1,2]. We filled a 150 mm diameter glass Petri dish with 60 mL of a 0.7 wt % aqueous slurry of nanocellulose. A smooth sheet of nanocellulose can be obtained by allowing the water to evaporate over the course of 2 – 3 days at ambient temperature (T ~ 25 °C). We found that the process can be accelerated if the slurry is placed in a 65 °C oven. The water evaporates within 2 hours instead of 2 – 3 days. However, the rapid drying results in a wrinkled nanocellulose sheet. These wrinkles can be smoothed by refilling the Petri dish with 60 mL of ultrapure water. The water evaporated from the petri dish in about 12 hours at ambient temperatures of 25 °C, reducing the time to fabricate wrinkle-free nanocellulose paper to ~ 14 hours (i.e. overnight).

### 3.2 Cleaning of substrates

**Nanocellulose paper and artist grade tracing paper:** Out of an abundance of caution, we exhaustively cleaned the paper substrates to remove adventitious soluble hydrophilic and hydrophobic materials. Control experiments using tracing paper as purchased without any cleaning showed no discernible difference in the yields or sizes of GUVs. Working in a chemical fume hood, we soaked the substrates in 100 mL of chloroform in a 500 mL glass beaker while applying occasional manual agitation. We allowed the substrate to soak for 30 minutes and then discarded the chloroform. We repeated the process with fresh chloroform. We removed the substrates from the chloroform and allowed residual solvent to evaporate by leaving the paper in the fume hood for 30 minutes. We then soaked the substrates in ultrapure water. After 30 minutes, we discarded the ultrapure water and repeated the process with fresh ultrapure water. We placed the papers flat in a glass petri dish before placing the dish in a 65 °C oven for 2 hours to dry.

**Regenerated Cellulose Dialysis Membranes:** We followed the manufacturer's instructions to remove the humectant glycerol and traces of heavy metals and sulfur compounds from the regenerated cellulose dialysis membrane. Briefly, we cut the dry dialysis tubing into 10 cm long pieces and agitate the tubing in 50 mL of ultrapure for 30 minutes. We then cut the tubing open lengthwise and transfer the tubing to new clean glass beaker filled with fresh ultrapure water for a further 60-minute incubation. We next agitate the tubing in a 10 mM solution of sodium bicarbonate at 80°C for 30 minutes. Then we incubated the tubing in a solution of 10 mM EDTA for 30 minutes at room temperature. We finally rinse the tubing under flowing ultrapure water for 10 minutes and transfer the tubing into a clean glass jar with ultrapure water at 80°C while stirring with a magnetic stirrer for 30 minutes. We removed the tubing pieces from the water and wound the wet membrane around a glass slide and allowed it to dry ambiently. Winding the tubing around a glass slide prevented excessive wrinkling or shrinking of the tubing as it dried.

**Glass slides and ITO-covered slides:** The electric field degrades ITO-covered slides each time it is used[3]. The degradation results in lower yields of GUVs and results in GUVs of smaller sizes[3]. Annealing in air at 150 °C for 20 minutes is reported to reverse these effects[3]. To ensure the highest possible yields from electroformation while nominally mimicking the pattern of use of this expensive substrate in a typical laboratory, we reuse pristine ITO-covered slides a maximum of five times. Before each use, we cleaned the slides to remove adventitious hydrophobic or hydrophilic materials by sonicating sequentially for 10 minutes in acetone, ethanol, and ultrapure water. We then dried the slides and annealed the slides in air at 150 °C for 20 minutes[3].

### 3.3 Hydrophobic modification of the substrates

**Silanization of glass slides:** We performed vapor phase silanization of plasma-cleaned glass slides using methyltrichlorosilane following previously reported protocols[4]. We placed 0.5 mL of neat methyltrichlorosilane in a 10 mL glass vial. Working quickly to minimize exposure to atmospheric moisture, we placed the uncapped glass vial containing the silane in a laboratory vacuum chamber with the plasma-cleaned glass slides. We also placed a Petri dish filled with the desiccant DrieRite$^{TM}$ in the vacuum chamber to further reduce ambient moisture. We allowed the reaction to proceed under vacuum overnight. The methyltrichlorosilane reacts with the silanol groups on the glass surface and covalently grafts methyl groups onto the glass thus rendering the surface hydrophobic[4].

**Silanization of tracing paper:** We perform solution phase silanization of pieces of tracing paper following previously reported protocols[5]. Working in a chemical fume hood, we prepared 10 mL of a 80/20 by volume mixture of neat anhydrous toluene and neat methyltrichlorosilane in a clean glass vial. We placed 9 pieces of tracing paper 9.5 mm in diameter into the silane solution, capped the vial, and allowed the reaction to proceed for 30 minutes. We then removed the paper from the silane solution and washed the paper with excess neat toluene 5 times to remove unreacted silanes. We allowed the paper to dry in the fume hood for 30 minutes, and then transferred the paper to a 65 °C oven for 2 hours.

### 3.4 Characterization of the substrates

**Scanning electron microscopy (SEM):** SEM images of the dry substrates were obtained using a field emission scanning electron microscope (GeminiSEM 500, Zeiss, Germany). The substrates were cut into small 2 × 2 mm squares and mounted on aluminum stubs using double-sided copper tape. A piece of copper tape was placed on top of the substrate at one edge and was connected to the stub to create a conduction path to minimize charging at the surface. The beam accelerating voltage was set to 1 kV. We used an Everhart-Thornley secondary electron detector to collect the secondary electrons that scattered from the surface. Images were captured at a lateral pixel resolution of 1.09 µm/pixel [1120 µm × 840 µm (1024 pixels × 768 pixels)], and 21 nm/pixel [22.76 µm × 17.07 µm (1024 pixels x 768 pixels)].

**Measurement of the average fiber diameter and length:** Using ImageJ, we measure the diameters of 82 randomly selected nanocellulose fibers from SEM images of the paper. The average diameter of the fibers was 34 ± 11 nm. To determine the nanofibril length, we measure the end-to-end length of 47 fibers. The average length of the fibers to be 5 ± 2 µm.

### 3.5 Growth of the GUVs

**Lipid solutions:** We dissolved DOPC powder in neat chloroform to obtain a stock solution at a concentration of 25 mg/mL. A lipid mixture consisting of 99.5:0.5 mol % DOPC: TopFluor®-Cholesterol was mixed to obtain a stock concentration of 10 mg/mL lipid. We prepared working solutions at a concentration of 1 mg/mL from this stock solution. All stock and working solutions were stored in a -20 °C freezer in Teflon-capped glass vials that were purged with argon.

**Deposition of Lipids:** We punched out circular disks with a diameter of 9.5 mm from the cleaned nanocellulose paper, tracing paper, and regenerated cellulose dialysis membrane, using a circle hole punch (EK Tools Circle Punch, 3/8 in.). We deposited 10 µL of the working solution of lipid using a glass syringe (Hamilton) onto the paper. Similarly, for gentle hydration and electroformation we spread 10 µL of the working lipid solution onto a 9.5 mm diameter circular area on the substrates. We used a 9.5 mm diameter paper cutout placed on the back surface of the slides to serve as a guide for spreading the lipid. All substrates were placed into a standard laboratory vacuum desiccator for a minimum of 1 hour to remove traces of solvent before proceeding to the growth stage.

### 3.6 Procedures for growth

**Gentle hydration on nanocellulose paper, tracing paper, and regenerated cellulose dialysis membranes:** We placed the dry solvent-free lipid-coated substrates into individual wells in a 48-

well plate. With a pipette, we added 150 µL of the hydrating solution at the bottom corner of the well to fully immerse the substrate. The 48-well plate was covered with a lid and the lipid-coated substrates were allowed to incubate in the growth buffer for 2 hours.

**Gentle hydration on glass:** We affixed circular PDMS gaskets (inner diameter × height = 12 × 1 mm) to construct a barrier around the dry solvent-free lipid films. We added 150 µL of the hydrating solution into the gaskets and carefully constructed chambers by placing a glass coverslip on the top surface of the PDMS gasket. We allowed the film to hydrate for 2 hours.

**Electroformation:** We used established protocols to prepare GUVs[3,6,7]. We affixed circular PDMS gaskets (inner diameter × height = 12 × 1 mm) to construct a barrier around the lipid film. We added 150 µL of the hydrating solution and used a second ITO-coated glass slide to form closed chambers. The ITO surfaces were connected to the leads of a function generator (33120A Agilent) with conductive copper tape. We applied a sinusoidal AC field at a field strength of 1.5 V/mm peak-to-peak and frequency of 10 Hz for 2 hours.

**Procedures for harvesting:** For methods that use a PDMS ring and a top slide or coverslip (gentle hydration, electroformation) we disassembled the chamber by carefully removing the top cover. For techniques that used the 48-well plate (gentle hydration on nanopaper, tracing paper, and regenerated cellulose membranes) no disassembly was required. We harvested the GUVs from each technique identically by gently aspirating 100 µL of the hydrating solution into a cut 1000 µL pipette tip. This procedure was repeated exactly 6 times on different regions to cover the whole substrate. On the seventh time, we aspirated all the liquid ~ 150 µL containing the GUVs and transferred the liquid into an Eppendorf tube. Aliquots were taken immediately for imaging.

**3.7 Quantification of the sizes and molar yields of the GUVs**

**Confocal microscopy of harvested vesicles:** We constructed imaging chambers by covalently bonding custom-made PDMS gaskets with a square opening (width × length × height = 6 × 6 × 1 mm) to glass microscope slides. Before use, we passivated the chamber with a solution of 1 mg/mL casein to prevent rupture of the GUVs on the bare glass[8]. We filled the passivated chamber with 58 μL of a 100 mM solution of glucose and added a 2 μL aliquot of the suspension of harvested GUVs. We allowed the GUVs to sediment for 3 hours before imaging. We captured images using an upright confocal laser-scanning microscope (LSM 880, Axio Imager.Z2m, Zeiss, Germany). We excited the TopFluor® dye with a 488 nm argon laser and collected fluorescence using a 10× Plan-Apochromat objective with a numerical aperture of 0.45. We imaged the entire area of the chamber using an automated tile scan routine (64 images [850.19 μm × 850.19 μm (3212 pixels × 3212 pixels)]). The routine used an autofocus feature at each tile location. The pinhole was set at 12.66 Airy Units which gave a confocal slice thickness of 80 μm.

**Image processing and data analysis:** We analyzed the confocal tilescan images using a custom MATLAB routine[9] (Mathworks Inc., Natick, MA). The routine thresholded the images and then applied a watershed algorithm to segment the fluorescent objects from the background. We used the native *regionprops* function to obtain the equivalent diameters and mean intensities of the segmented objects. GUVs were selected from the detected objects based on the coefficient of variance of the intensities. GUVs fall within 1 – 2 times the full width at half the maximum (FWHM) of the highest peak in the coefficient of variance histogram. Once selected, we used MATLAB native routines to obtain the diameters and the counts of the GUVs. We plot size histograms, calculate the molar yields, and perform statistical tests.

**Size distributions:** We used MATLAB to plot histograms of the vesicle diameters for each repeat using 1 μm wide bins. The bins were normalized to show the GUV counts per μg of lipid deposited

(10 μg for all substrates). We take a mean of the counts in each bin from the histograms of the 5 independent repeats per surface to plot the results in Supplementary Fig. 2 and 3a.

**Calculation of median sizes:** To determine the median GUV size for each surface, we calculated the median diameter of the population from each of the 5 independent repeats. We then take a mean of these 5 values to prepare the plot in Supplementary Fig. 3b.

**Calculation of extreme sizes:** To determine the extreme sizes in each population we calculated the mean of the largest 100 vesicles for each of the 5 independent repeats. We then take a mean of these 5 values to prepare the plot in Supplementary Fig. 3c.

**Calculation of molar yield:** We obtain the moles of lipid in *N* GUVs from a harvested suspension using Equation (3.7.1).

$$mol_{GUV} = \frac{2\pi V_h}{N_A\, A_{hg} V_{al}} \sum_{i=1}^{N} (d_i)^2 \tag{3.7.1}$$

In this equation, $N_A$ is Avagadro's number, $A_{hg}$ is the lipid headgroup area, $V_h$ is the volume of the harvested suspension, $V_{al}$ is the volume of the aliquot, and $d_i$ is the diameter of vesicle *i*. The factor of 2 accounts for the 2 lipid leaflets in a bilayer. The molar amount of GUVs of a given diameter can be obtained by summing GUVs of a specific diameter. In the main text, we defined three size ranges, small GUVs ($1 \le d_i < 10$), large GUVs ($10 \le d_i < 50$) and very large GUVs ($d_i \ge 50$).

$$mol_{GUV} = mol_{\text{small GUVs}} + mol_{\text{large GUVs}} + mol_{GUVs \geq 50} \tag{3.7.2}$$

$$mol_{GUV} = \frac{2\pi V_h}{N_A\, A_{hg} V_{al}} \left( \sum_{i=1}^{N_{d<10}} (d_i)^2 + \sum_{i=d\geq 10}^{N_{d<50}} (d_i)^2 + \sum_{i=d\geq 50}^{N} (d_i)^2 \right) \tag{3.7.3}$$

To obtain the molar yield, $Y_{mol}$ we divide the mols of lipids in the GUV membranes with the total mols of lipid deposited on the substrate (kept the same for all substrates)and report the ratio as a percentage.

$$Y_{mol} = \frac{mol_{GUV}}{mol_{tot}} \times 100\% \quad (3.7.4)$$

**Statistical Tests:** We performed all statistical tests in MATLAB. For statistical testing, the groups were assigned by technique, group 1: gentle hydration on glass, group 2: gentle hydration on regenerated cellulose dialysis membranes, group 3: gentle hydration on nanopaper, group 4: gentle hydration on tracing paper, group 5: gentle hydration on silanized glass, group 6: gentle hydration on silanized tracing paper, group 7: electroformation. We test for statistical significance of differences in the mean molar yields by performing a balanced one-way Analysis of Variance (ANOVA) followed by a post-hoc Tukey's honestly significant test (HSD). An ANOVA assumes that the repeats drawn for each group comes from a normal distribution and that the variances between each group is equal. We conducted an Anderson Darling test to determine normality. The results of the test are shown in Supplementary Table 2. All repeats within a group were consistent with being drawn from a normal distribution. We conducted a Bartlett's test to determine if the variances of the groups were equal. Results of the tests are shown in Supplementary Table 2. The variances between groups were equal. Thus, the data satisfies the criteria for an ANOVA. We show the ANOVA table and post-hoc HSD tables in Supplementary Table 3. We also make summary conclusions in the table.

### 3.8 In situ imaging on the surface of the substrate

**Imaging on the surface:** We prepared the lipid-coated substates following the same standard procedures we used for quantification of the molar yield. Instead of using the 48-well plates for growth, we used circular PDMS gaskets (inner diameter × height = 12 mm × 1 mm) affixed to glass slides. After hydrating with 150 µL of growth buffer, we sealed the chamber with a glass coverslip. We captured images after 2 hours of growth using a confocal microscope. We excited the TopFluor® dye with a 488 nm argon laser and imaged using a 10× Plan-Apochromat objective with a numerical aperture of 0.45.

#### 3.9 Fluorescence recovery after photobleaching vesicle buds and isolated vesicles

**Vesicle buds:** Fluorescence recovery after photobleaching (FRAP) experiments were performed using a 488 nm argon laser. Circular regions of closely-packed vesicle buds (~40 µm in diameter) were bleached at 100 % laser power. Time lapse images of the subsequent recovery were collected at 1 second intervals for 250 seconds using 0.1 % of the laser power. The mean intensity of the region of interest was measured using ImageJ. The plots were normalized to the pre-bleach intensity which was set to 1.

**Isolated vesicles:** To obtain closely-packed isolated vesicles, we pooled vesicles from 3 tracing paper samples to obtain 450 µL of the vesicle solution. We centrifuged the vesicles at 10,000 × g for 3 minutes and remove the top 300 µL leaving behind a 150 µL solution of concentrated vesicles. A 30 µL aliquot of the concentrated vesicles was added to a PDMS chamber containing 30 µL of glucose, and the chamber was sealed with a glass coverslip. We allowed the vesicles to sediment for 5 hours before imaging. We performed FRAP experiments on the isolated vesicles using the same conditions as those used for the vesicle buds.

#### 3.10 Time lapse images of vesicle buds

We used time-lapse imaging to capture the dynamics of the vesicle buds on the surface of the substrates. We excited the dye in the vesicles using a 488 nm argon laser and collected the fluorescence using a 20× water-dipping Plan-Apochromat objective with a numerical aperture of 1.0. We captured images of the vesicle buds on the surface of the tracing paper at 5 second intervals 3 minutes after hydration. Typical images covered an area of approximately 200 μm × 200 μm and had a pixel resolution of 0.12 μm.

#### 3.11 Scale up and calculation of cost

**Scale up:** To carry out scale-up experiments we used a Paasche airbrush and compressor to spray lipid onto whole sheets of tracing paper 9" × 12" in size. We used PVA-coated gloves when working with the chloroform and performed all experiments in a chemical fume hood. To coat the paper, we prepared 9.6 mL of 1 mg/mL DOPC:TFC (99.5:0.5 mol %) lipid mixture in chloroform to mimic the same lipid mass as the 48-well plate experiments. We used a 0.66 mm cap at the end airbrush, set the compressor pressure to 20 PSI, and brushed the lipid across the surface at a rate of approximately 1" per second. We placed the lipid-coated sheet under vacuum for 1 hour to allow residual solvent to evaporate, and then placed the sheet in a 9" × 13" Teflon coated baking pan. We hydrated the sheet in 100 mL of 100 mM solution of sucrose and allowed the GUVs to grow for 1 hour. To harvest the GUVs, we pipette across the surface using a 1 mL plastic transfer pipette. We store the harvested GUVs in 50 mL Eppendorf tubes.

**Calculation of cost:** The total fixed material costs of fabricating GUVs include the cost of the substrate and the cost of the lipids. To obtain the substrate costs, we calculated the cost per unit area ($mm^2$) of each substrate using the lowest prices posted in July 2020 on the websites of large multinational suppliers of scientific materials (Supplementary Table 4). To determine the number of vesicles harvested from a given area of substrate, we normalized the size histograms by dividing the counts in each bin (bin widths of 1 μm) by the area ($mm^2$) of substrate that was used to grow vesicles, which for all of the experiments was 70.88 $mm^2$. The counts per area for each bin was divided by the substrate cost per area to obtain the substrate cost per vesicle (Fig. 5a). Additionally, all methods require the use of solvent, buffers, and fluid receptacles. For electroformation, additional equipment such as a function generator and a source of electrical power is required for directing the formation of vesicles. We do not consider these additional costs in our calculations.

**Cyclic reuse of tracing paper:** After harvesting the vesicles we rinsed the paper under a flowing stream of ultrapure water for 30 seconds and then soaked the paper in 10 mL of ultrapure water for 30 minutes. We occasionally agitated the paper in the water by manually shaking the container. This procedure removes any water-soluble residual sugars from the prior cycle of growth. We then moved the tracing paper to a 65°C oven to dry for 1 hour. To remove any residual lipid, we soaked the tracing paper in 10 mL of neat chloroform for 30 minutes. We occasionally agitated the paper in the chloroform by manually shaking the container. We then carefully removed the tracing paper and placed it in a laboratory vacuum chamber for 1 hour to remove traces of chloroform. After drying, the standard procedures for growth were employed to start the next cycle of growth.

### Supplementary Figures

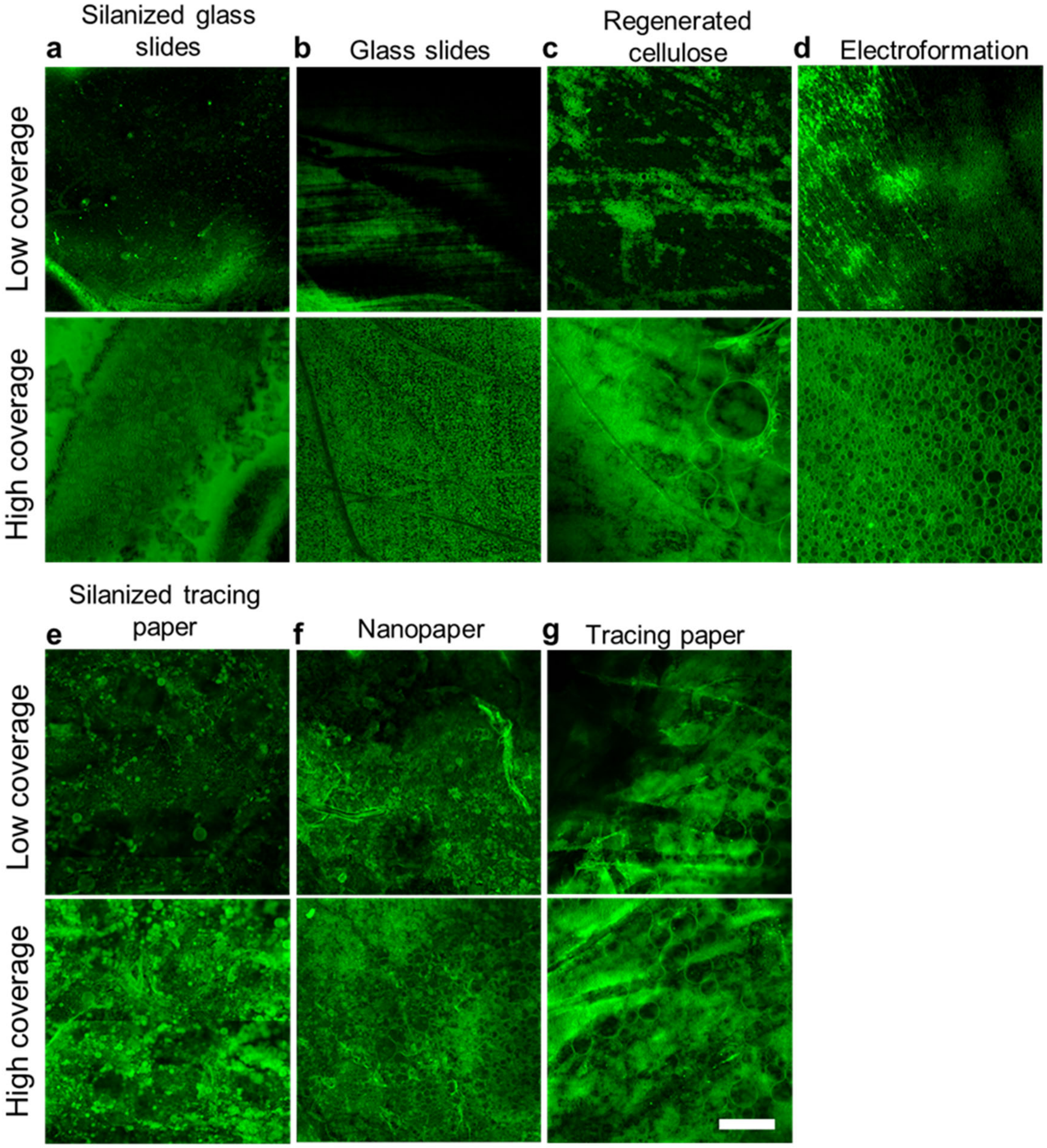


**Supplementary Figure 1.** Characterization of the coverage of vesicle buds on the substrates. **a-d** Confocal images of regions with low and high vesicle bud coverage on **a** silanized glass slides, **b** glass slides, **c** regenerated cellulose and **d** electroformation. **e-g** Images of regions with low and high vesicle coverage on **e** silanized tracing paper, **f** nanopaper, and **g** tracing paper. Scale bar 100 μm.

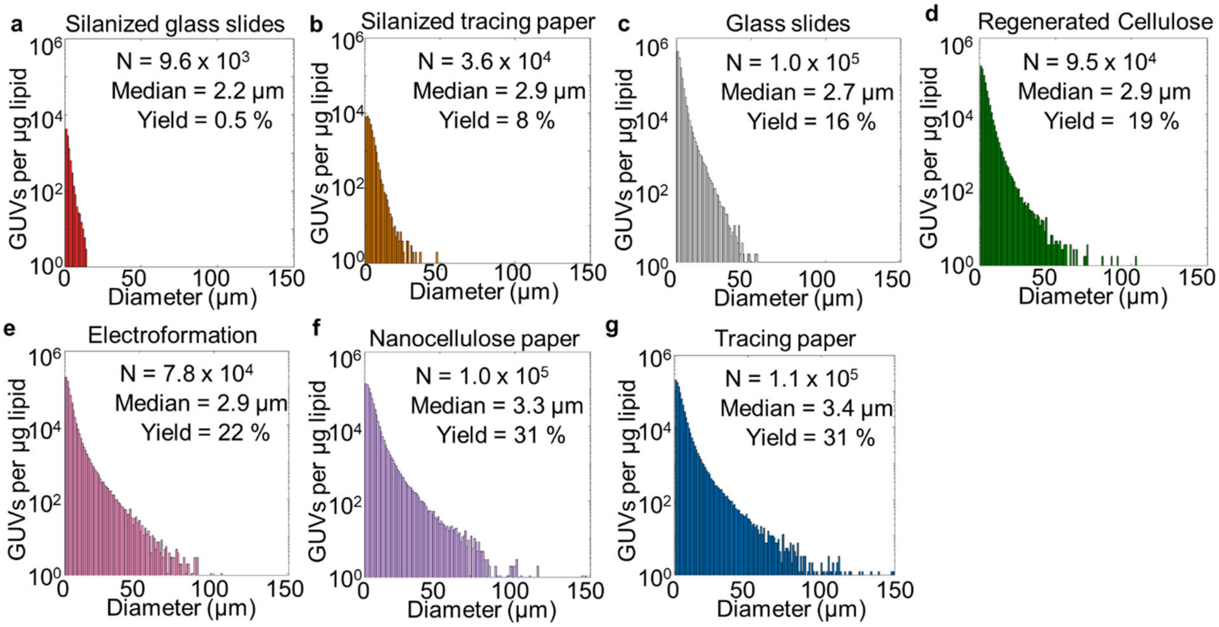


**Supplementary Figure 2.** Distribution of vesicle diameters that were obtained from the assorted substrates**.** Normalized histograms of the diameters of the GUVs harvested from **a** silanized glass slides, **b** silanized tracing paper, **c** glass slides, **d** regenerated cellulose dialysis membranes, **e** electroformation on ITO-covered glass slides, **f** nanocellulose paper, and **g** tracing paper. The number of GUVs counted, the median size, and the molar yield is added to each plot. Bin widths are 1 μm. Note the logarithmic scale on the *y* axis.

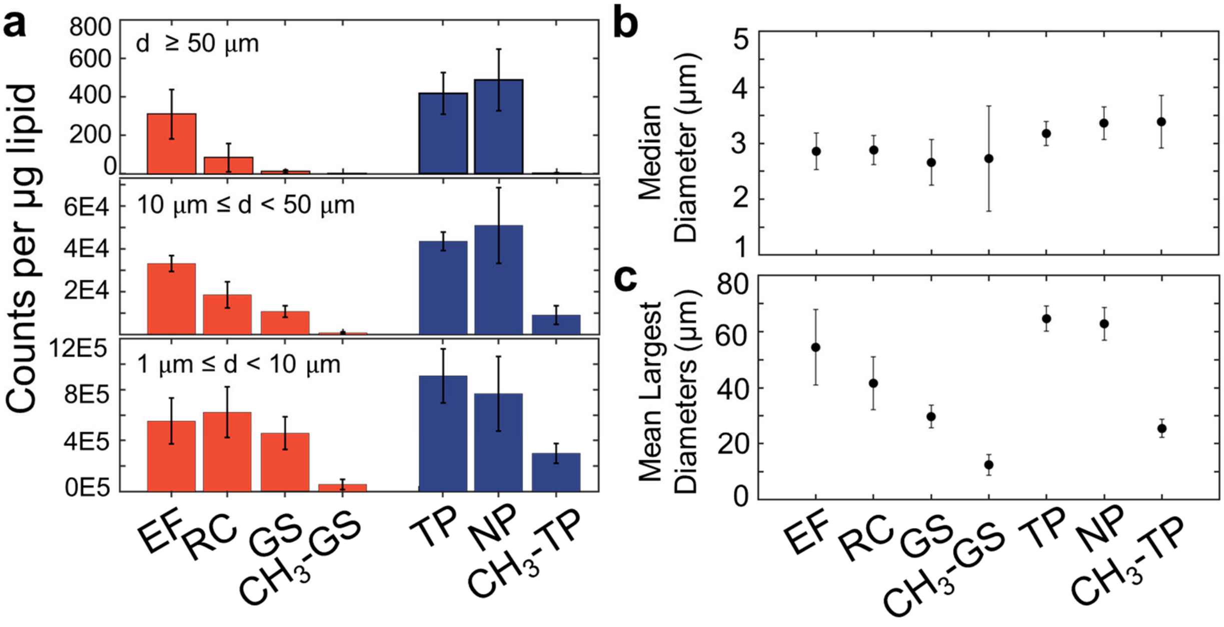


**Supplementary Figure 3.** Summary statistics of the GUVs obtained from the assorted surfaces. **a** Mean GUV counts normalized by the amount of lipid deposited for each surface. Each plot shows the mean counts within the different diameter ranges specified. Counts of GUVs larger than or equal to 50 µm in diameter (top plot) for electroformation on ITO-covered glass slides (EF), regenerated cellulose dialysis membranes (RC), glass slides (GS), silanized glass slides ($CH_3$-GS), tracing paper (TP), nanocellulose paper (NP) and silanized tracing paper ($CH_3$-TP) were 309 ± 129, 83 ± 74, 11 ± 8, 0, 467 ± 134, 488 ± 160, 1.5 ± 0.7. In the same order, counts of vesicles greater than or equal to 10 µm and less than 50 µm (middle plot) were 3.29E4 ± 3.70E3, 1.85E4 ± 6.10E3, 1.08E4 ± 2.67E3, 740 ± 503, 4.94E4 ± 1.42E3, 5.09E4 ± 1.77E3, 0.91E4 ± 4.31E3 and counts of vesicles greater than or equal, to 1 µm and less than 10 µm (bottom plot) were 5.54E5 ± 1.81E4, 6.23E5 ± 2.00E4, 4.54E5 ± 1.30E4, 0.56E5 ± 4.01E4, 9.08E5 ± 2.12E4, 7.68E5 ± 2.92E4, 2.98E5 ± 7.72E4. **b** Scatter plot showing the median diameter of the GUV populations obtained from the surfaces. **c** Scatter plot of the mean diameter of the largest 100 vesicles for each of the surfaces. The mean median diameters from the 5 repeats for EF, RC, GS , $CH_3$-GS, TP, NP, and $CH_3$-GS were 2.8 µm ± 0.3 µm, 2.9 µm ± 0.3 µm, 2.7 µm ± 0.4 µm, 2.7 µm ± 0.9 µm, 3.2 µm ± 0.2 µm, 3.4 µm ± 0.3 µm, and 3.4 µm ± 0.5 µm respectively. In the same order, the mean diameters of the largest 100 vesicles were 54 µm ± 14 µm, 42 µm ± 9 µm, 30 µm ± 4 µm, 12 µm ± 4 µm, 65 µm ± 5 µm, 63 µm ± 6 µm, and 26 µm ± 3 µm. Error bars are standard deviations from the mean ($N = 5$).

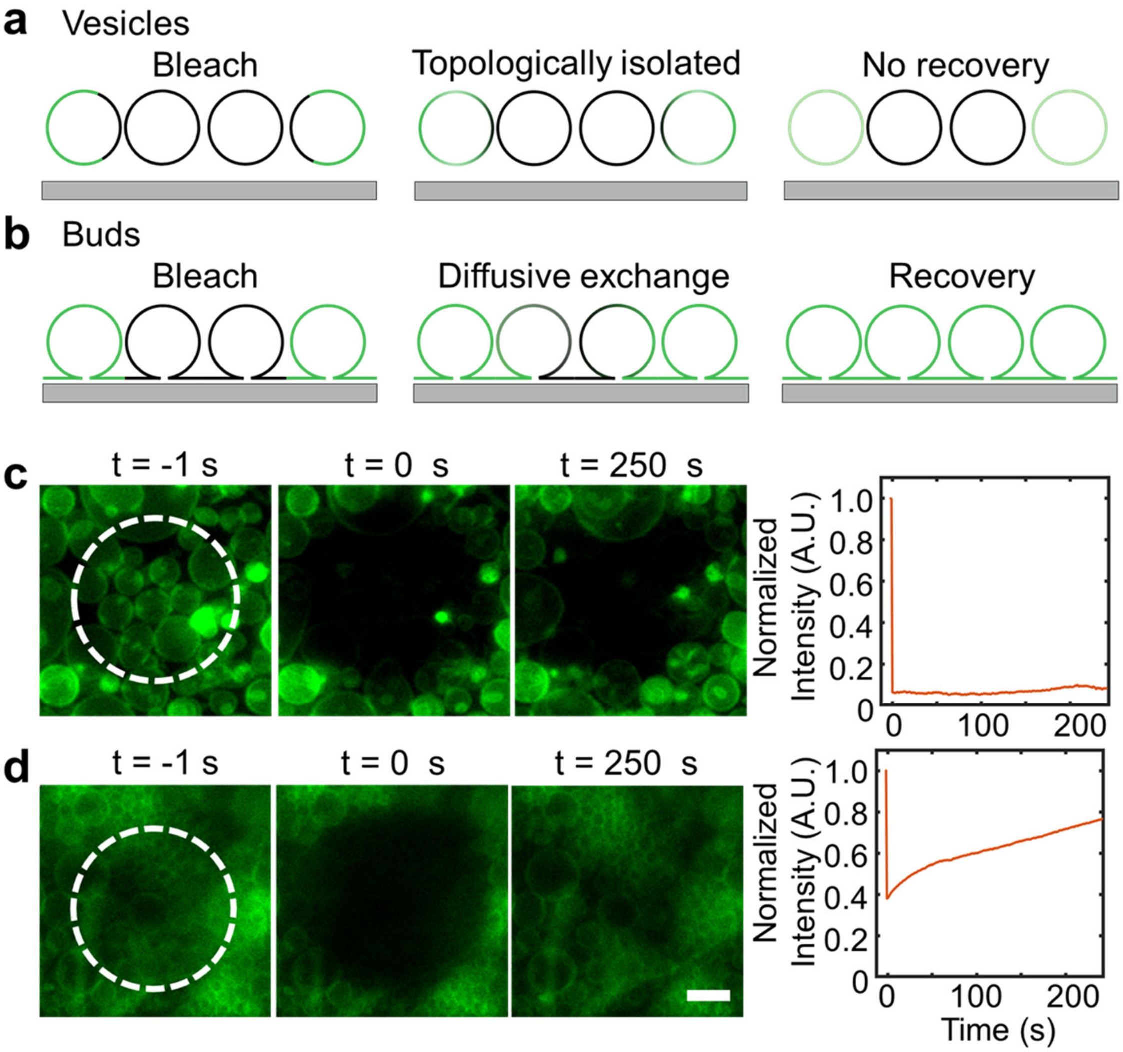


**Supplementary Figure 4.** Objects on the surfaces are connected vesicular buds and not topologically isolated closed vesicles. **a** Cross sectional schematic of the expected behavior after photobleaching of the fluorophore in the membranes of a topologically isolated closed vesicles and **b** connected vesicle buds. The black lines indicate photobleached membranes. Stills from FRAP experiments showing the frame before application of a circular bleach pulse, the first frame after application of the bleach pulse, and the final frame in the experiment for **c** isolated vesicles and **d** vesicle buds on the surface of tracing paper. The dotted white circles in the first panels shows the region that was photobleached. The plots show normalized FRAP recovery curves. Scale bar 10 µm.

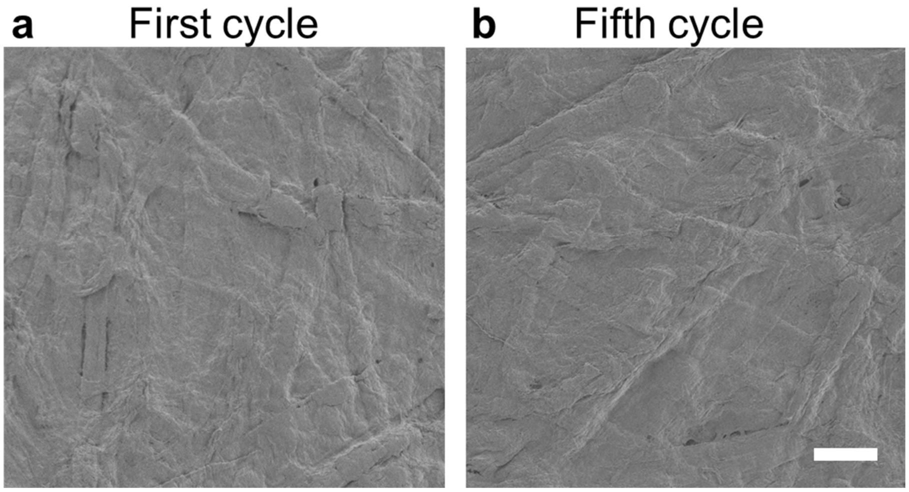


**Supplementary Figure 5.** Characterization of the reused tracing paper surface. SEM images showing the surface structure of the tracing paper remains unchanged. **a** image after the first growth cycle and **b** image after the fifth growth cycle. Scale bar 50 µm.

### Supplementary Tables

| | Value | Electro. | Regen. Cellulose | Glass | Silanized Glass | Tracing Paper | Nanopaper | Silanzied Tracing Paper |
|---|---|---|---|---|---|---|---|---|
| Anderson Darling | p | 0.5241 | 0.3831 | 0.847 | 0.4885 | 0.5251 | 0.3119 | 0.3739 |
| Bartlett's Test | p | 0.3462 | | | | | | |

**Supplementary Table 1:** Results of the Anderson-Darling and Bartlett tests. NS not significant, * $p < 0.05$, ** $p < 0.01$ *** $p < 0.001$.

| Source | SS | df | MS | F | Prob > F, (p-value) |
|---|---|---|---|---|---|
| Columns | 3725.24 | 6 | 620.873 | 90.53 | 5.0591E-17 |
| Error | 192.04 | 28 | 6.858 | | |
| Total | 3917.27 | 34 | | | |

| Group 1 | Group 2 | p-value | Significance | Comments |
|---|---|---|---|---|
| Glass | Electro-formation | 0.00542 | ** | Effect of electric field significant |
| Glass | Regenerated Cellulose | 0.383 | NS | Effect of permeability not significant |
| Glass | Nanopaper | 5.28E-08 | *** | Effect of curvature significant |
| Glass | Tracing Paper | 4.31E-08 | *** | Effect of curvature significant |
| Glass | Silanized Glass | 6.23E-08 | *** | Effect of hydrophilicity significant |
| Glass | Silanized TP | 0.00304 | ** | Effect of hydrophilicity significant |
| Electro-formation | Nanopaper | 5. 22E-04 | *** | Effect of curvature significant |
| Electro-formation | Tracing Paper | 1.72E-04 | *** | Effect of curvature significant |
| Electro-formation | Regenerated Cellulose | 0.439 | NS | Effect of electric field not significant |
| Electro-formation | Silanized Glass | 3.71E-08 | *** | Effect of hydrophilicity significant |
| Electro-formation | Silanized TP | 1.14E-07 | *** | Effect of hydrophilicity significant |
| Tracing Paper | Nanopaper | 0.999 | NS | Effect of manufacturing paper not significant |
| Tracing Paper | Regenerated Cellulose | 9.63E-07 | *** | Effect of curvature significant |
| Tracing Paper | Silanized Glass | 3.71E-08 | *** | Effect of curvature significant |
| Tracing Paper | Silanized TP | 3.71E-08 | *** | Effect of hydrophilicity significant |
| Nanopaper | Regenerated Cellulose | 2.71E-06 | *** | Effect of curvature significant |
| Nanopaper | Silanized Glass | 3.71E-08 | *** | Effect of curvature significant |
| Nanopaper | Silanzied TP | 3.71E-08 | *** | Effect of hydrophilicity significant |

| | | | | |
|---|---|---|---|---|
| Regenerated Cellulose | Silanized Glass | 3.73E-08 | *** | Effect of hydrophilicity significant |
| Regenerated Cellulose | Silanized TP | 1.18E-05 | *** | Effect of hydrophilicity significant |
| Silanized TP | Silanized Glass | 0.00162 | ** | Effect of curvature significant |

**Supplementary Table 2:** ANOVA table and table of p-values from post hoc Tukey's HSD tests of the molar yields of GUVs obtained from gentle hydration on glass, regenerated cellulose, nanocellulose paper, tracing paper, silanized glass, and silanized tracing paper and from electroformation on ITO slides.

| Parameter | Description | Magnitude | Source |
| --- | --- | --- | --- |
| $\kappa_a$ | Area expansion modulus | $80 - 200 \times 10^{-3}$ J m$^{-2}$ | [10] |
| $\xi$ | Adhesion energy | $-0.1$ to $-50 \times 10^{-8}$ J m$^{-2}$ | [10–12] |
| $\lambda$ | Membrane edge energy | $1 \times 10^{-11}$ J m$^{-1}$ | [10] |
| $\kappa_G$ | Gaussian bending modulus | $-0.9\kappa_B$ to $-1.0\kappa_B$ | [13] |
| $\kappa_B$ | Bending modulus | $8.22$ to $82.2 \times 10^{-20}$ J | [10] |

**Supplementary Table 3:** Values for the physical parameters of phosphocholine membranes reported in the literature.

| Substrate | Cost (USD) | Unit of measure | Cost (USD) per $mm^2$ | Source of cost |
|---|---|---|---|---|
| **Glass Slide** | 733.61 | 1440 glass slides, 25 mm × 75 mm | 2.72E-04 | Fischer Scientific |
| **ITO-slide** | 144 | 10 slides, 25 mm × 25 mm | 2.30E-02 | Sigma Aldrich |
| **Regenerated Cellulose** | 224.5 | 1 roll, 30,000 mm × 90 mm | 8.31E-05 | Fischer Scientific |
| **Tracing paper** | 26.93 | 500 sheets, 228.6 mm × 304.8 mm | 7.73E-7 | Amazon |

**Supplementary Table 4**: Material costs of substrates used to assemble GUVs. Costs per $mm^2$ were calculated using the most economical option available online at various large scientific supply companies.
